# Lamin A/C maintains genome topology and regulates transcriptional programs essential for virus-driven B cell activation

**DOI:** 10.64898/2026.01.12.699161

**Authors:** Lisa B Caruso, Davide Maestri, Andrew Kossenkov, Aaron R. Goldman, Joel Cassel, Samantha Soldan, Paul M. Lieberman, Italo Tempera

## Abstract

Lamin A/C is a crucial structural component of the nuclear lamina that influences chromatin organization and gene regulation. In this study, we demonstrate that lamin A/C is vital for maintaining higher-order genome organization and transcriptional programs that support EBV-driven B-cell activation. Loss of lamin A/C in a B-lymphoblastoid cell line caused significant three-dimensional reorganization of the genome, evidenced by the loss of long-range chromatin loops, an increase in short-range contacts, and redistribution of H3K9me2- marked heterochromatin. These structural disruptions were linked to widespread changes in gene expression affecting metabolic, signaling, and differentiation pathways. Mechanistically, lamin A/C influences the nuclear positioning and transcription of CTCF-bound loci by preventing their relocation to the periphery and their association with lamin B1. Blocking H3K9me2 deposition mimicked the transcriptional effects of lamin A/C depletion and revealed increased sensitivity to PI3K inhibitors. Overall, our results identify lamin A/C as a key organizer of genome structure and epigenetic regulation in EBV-infected B cells, uncovering a lamin-dependent pathway that connects nuclear architecture, metabolism, and viral disease processes.

## INTRODUCTION

Epstein-Barr virus (EBV), a member of the gamma herpesvirus subfamily, was the first human oncogenic herpesvirus to be identified and characterized (1,2). It is a highly successful pathogen, infecting approximately 95% of the global adult population and establishing a persistent lifelong infection (3). Persistent EBV infection is usually asymptomatic but can be associated with increased risk of several malignancies, particularly in immunocompromised individuals and specific geographic and ethnic groups, contributing to approximately 5% of all cancer cases globally (4). EBV’s ability to establish lifelong persistence depends on its capacity to hijack host cellular programs, particularly those governing B-cell differentiation and proliferation (5,6). In latently infected B cells, EBV expresses a restricted set of viral genes that facilitate immune evasion, maintain latency, and promote oncogenic transformation (7). EBV can activate different gene expression programs, referred to as latency type I, II, and III, leading to the expression of specific viral proteins and RNAs (8). The viral genes expressed during latent infection strongly depend on the host cell and the tumor type (6). To modulate B-cell fate and activation (9,10), EBV-encoded proteins engage in host regulatory pathways, including those involved in epigenetic control (11–13). Recent studies have highlighted the critical role of epigenetic modifications in sustaining EBV latency and virus-driven proliferation. Key viral effectors such as EBNA2, EBNA-LP, the EBNA3 family, and LMP1 (14–24) co-opt host epigenetic regulators—including DNMT1, EZH2, YY1, KDMs, and others—to reprogram the host epigenome and drive a transcriptional landscape conducive to viral persistence and oncogenesis (16,24–29). Our group and others have demonstrated that EBV infection also reshapes the host genome’s three-dimensional (3D) architecture to fine-tune the expression of cellular genes (29–33). EBV latent proteins have been shown to modulate, both directly and indirectly, the activity of key chromatin architectural regulators, including CTCF, cohesin, PARP1, and YY1, facilitating the virus’s ability to reprogram genome organization (34–39). These findings suggest that EBV regulates host gene expression through an intricate network involving histone modifications, 3D chromatin architecture, and epigenetic regulators. However, the functional interaction of these distinct epigenetic mechanisms to coordinate gene regulation during EBV latency remains poorly understood.

In recent years, the interaction between chromatin and the nuclear lamina has emerged as a critical mechanism for regulating the deposition of specific histone modifications and the formation of chromatin loops across genomic regions associated with the nuclear periphery (40–44). The nuclear lamina is a complex protein network located mainly beneath the inner nuclear membrane (45,46). This strategic localization is essential for creating a functional interface between the nuclear envelope and the genome (47). The lamina plays a crucial role in maintaining nuclear architecture, organizing chromatin, and regulating gene expression. The primary components of the nuclear lamina are type V intermediate filament proteins known as lamins, which are broadly classified into A-type lamins (lamins A and C) and B-type lamins (lamins B1 and B2). B-type lamins are universally expressed across all cell types and are vital for nuclear structure, mechanical integrity, and embryonic development. They are primarily found at the nuclear periphery, where they associate mainly with transcriptionally inactive heterochromatin, contributing to chromatin anchoring and gene silencing (48). In contrast, A-type lamins are expressed in a tissue-specific and developmentally regulated manner. They are involved in various cellular processes, including cellular senescence, lineage differentiation, and DNA damage repair. A-type lamins significantly enhance the mechanical stiffness of the nucleus, helping to maintain its shape and resist mechanical deformation (49). Unlike B-type lamins, A-type lamins are not confined to the nuclear periphery; they are also present throughout the nucleoplasm, interacting with both heterochromatin and euchromatin (50). This broader localization allows A-type lamins to affect chromatin accessibility and 3D genome organization, linking nuclear structure to dynamic gene regulation (51). Fundamental to the role of lamin A/C in gene expression regulation is its interaction with DNA regulatory elements such as CTCF. While the exact mechanisms of how lamin A/C and CTCF collaborate are still being investigated, there is evidence that the two proteins can directly interact, possibly influencing each other’s function (52,53). Lamin A/C can impact the positioning and organization of chromatin (54), which in turn can affect CTCF binding sites and its ability to regulate gene expression (52,55). Both lamin A/C and CTCF are involved in shaping the 3D organization of the genome, with lamin A/C primarily influencing the structural framework and CTCF contributing to the formation of chromatin loops and topologically associating domains.

Our group has demonstrated that the 3D organization of the EBV epigenome is crucial in regulating viral gene expression and sustaining latent infection. Several epigenetic mechanisms influence the regulation and expression of EBV latency programs, including chromatin remodeling, histone modifications, and DNA methylation of the EBV genome (11). Collectively, our studies so far highlight that multiple epigenetic mechanisms contribute to EBV latency; however, the functional interaction of these mechanisms in coordinating viral gene regulation remains largely unresolved. In our previous work, we elucidated aspects of host-EBV virus interaction by focusing on the role of the host nuclear lamina in modulating viral gene expression and the epigenetic landscape during EBV latency. We demonstrated that EBV latent infection alters the composition of the host nuclear lamina and is associated with the expression of lamin A/C and the formation of lamin-associated domains (LADs) across the EBV genome (56). Our findings highlight a dynamic interaction between the EBV genome and nuclear lamins during latency. Notably, EBV infection leads to nuclear accumulation of lamin A/C, which binds to the viral genome and facilitates its repositioning toward the nuclear periphery. This relocalization is associated with reduced viral gene expression, suggesting a role for lamin A/C in the epigenetic silencing of the EBV genome. A similar role of lamin A/C in viral lytic reactivation was found for herpes simplex virus type 1 (HSV-1) (57). It was shown that A-type lamins are required for targeting the HSV genome to the nuclear periphery for assembling the early replication compartments of the infected cell nucleus and for preventing or reducing heterochromatin formation on the viral immediate-early lytic gene promoters. The same study also highlighted the pivotal role of lamin A/C in organizing chromatin remodeling and histone modification enzymes that regulate both euchromatin and heterochromatin (57). Whether the EBV-induced reorganization of the nuclear space affects the regulation of host gene expression remains unknown. This study aims to investigate whether EBV-induced modulation of lamin A/C expression contributes to transcriptional reprogramming in B cells. Specifically, our study focuses on understanding which host cellular pathways are impacted by altered lamin A/C expression and how EBV hijacks B-cell pathways to its biological advantage. To address our questions, we used next-generation sequencing techniques to assess how the deletion of lamin A/C in an EBV-infected lymphoblastoid cell line (LCL) influences gene expression and 3D chromatin structure. We found that the lack of Lamin A/C transcriptionally affects many pathways regulating B-cell metabolism and cell proliferation. Metabolism is a key player in regulating B-cell differentiation, development, and functions, including antibody production (58,59). Importantly, B cells possess the capacity to dynamically reprogram their metabolic pathways to accommodate the varying energetic and biosynthetic demands associated with different stages of their life cycle and during immune responses (60). Dysregulation of these metabolic processes has been implicated in the pathogenesis of various diseases, including B-cell lymphomas (61). EBV has evolved mechanisms to manipulate B-cell metabolism to create a cellular environment favorable for its own replication, persistence, and immune evasion. This metabolic reprogramming is a critical facet of EBV’s ability to establish latency and promote oncogenic transformation (62–65). Elucidating the role of lamin A/C in host gene regulation may provide new insights into how EBV establishes a cellular environment that supports viral latency and virus-driven B-cell proliferation. Understanding the interplay between nuclear lamina dynamics and host gene regulation in the context of EBV infection may uncover novel mechanisms of viral pathogenesis and identify potential novel therapeutic targets for EBV-associated diseases.

## MATERIAL AND METHODS

### Cell culture and treatment

Cell lines were maintained in a humidified atmosphere containing 5% CO_2_ at 37°C. EBV-positive B cells (lymphoblastoid cell line, LCL GM12878) were cultured in suspension in RPMI 1640 medium supplemented with fetal bovine serum at a concentration of 15% for type III latency. HEK 293T cells (human embryonic kidney cells) were cultured in adhesion in DMEM medium supplemented with fetal bovine serum at a concentration of 10%. All cell media were supplemented with 1% penicillin-streptomycin and plasmocin at a concentration 1:500 (Invivogen, ant-mpp). Treatment with the G9a inhibitor UNC0631 (Med Chem Express, HY-13808) was given 72 h before collection at a concentration of 1 μM. Treatment with acalisib (GS-9820) (Selleckchem, S5818) was given 72 h before collection at a concentration of 1 μM and 0.1 μM.

### CRISPR Cas9 mutagenesis

B cells were transduced with lentiviruses expressing sgRNAs as previously described (66). Briefly, lentiviruses were produced by transfection of 293T cells, which were plated in a 6-well dish at a density of 600,000 cells/well in 2 mL DMEM supplemented with 10% fetal bovine serum 24 hours before transfection. Transfection was carried out using the TransIT-LT1 Transfection Reagent (Mirus). Two solutions were prepared for each well: one solution with 4 mL of LT1 diluted in 16 mL of Opti-MEM (Corning, 31985070) and incubated at room temperature for 5 min. The second solution contained 150 ng pCMV-VSVG (Addgene, 8454), 400 ng psPAX2 (Addgene, 12260), and 500 ng p-lentiGuide-puro expression vector (Addgene, 52963), in Opti-MEM (final volume of 20 mL). The two solutions were then mixed and incubated at room temperature for 30 min, added dropwise to wells, and gently mixed. Plates were returned to a 37°C humidified chamber with 5% CO_2_. The following day, the transfection medium was replaced with RPMI 1640 with 10% fetal bovine serum. Virus supernatant was harvested 48- and 72h post-transfection, filtered through a 0.4 µM sterile filter, and added to target B cells. Target cell selection began 48 hours post-transduction by the addition of puromycin 3 µg/ml. sgRNA sequences used were as follows: Control: ATTTCGCAGATCATCGACAT, LMNA: AGTTTAAGGAGCTGAAAGCG.

### Western blot analysis

Cell lysates were prepared in radioimmunoprecipitation assay (RIPA) lysis buffer (50 mM Tris-HCl, pH 7.4, 150 mM NaCl, 0.25% deoxycholic acid, 1% NP-40, 1 mM EDTA; Millipore, 20-188) supplemented with protease inhibitor cocktail (ThermoFisher Scientific, A32955). Protein extracts were obtained by centrifugation at 3,000 × g for 10 min at 4°C. Protein concentration was measured using a bicinchoninic acid (BCA) protein assay (Pierce, 23227). Lysates were boiled with 2X Laemmli sample buffer (Bio-Rad, 161-0737) containing 2.5% 2-mercaptoethanol (Sigma-Aldrich, M3148-25ML). Proteins were resolved by gel electrophoresis on a 4 to 20% polyacrylamide gradient Mini-Protean TGX precast gel (Bio-Rad, 4561096) and transferred to an Immobilon-P membrane (Millipore, IPVH00010) using a Power Blotter XL System (Invitrogen). Membranes were blocked in 5% milk in PBS-T (PBS with 0.05% Tween-20) for 1 h at room temperature and incubated overnight at 4°C with primary antibodies against lamin A/C (Active Motif, 39287), lamin B1 (Abcam ab16048), β-actin (Cell Signaling 12262), CTCF (Active Motif 61311), H3 (Abcam, ab1791), H3K9me2 (Abcam, ab1220), Akt (Cell Signaling, 9272) and p-Akt (Invitrogen, PAKT-140AP) as per manufacturer’s recommendation. Membranes were washed 5 min 3 times, incubated for 1 h with the appropriate secondary antibody (goat anti-rabbit IgG-HRP, Jackson ImmunoResearch, 111-035-003 or rabbit anti-mouse IgG-HRP, Jackson ImmunoResearch, 315-035-003) at a dilution of 1:10,000. Membranes were then washed, and the signal was detected by enhanced chemiluminescence (SuperSignal West Dura Extended Duration Substrate ThermoFisher Scientific, 37071). Quantification was performed using iBright Analysis Software version 5.3.0 (ThermoFisher Scientific).

### RNA extraction and RNA-seq

Total RNA for lamin A/C knockout experiments was isolated from 2 × 10^6^ cells using a PureLink RNA Mini Kit (Invitrogen, 12183025) according to the manufacturer’s protocol. RNA samples were submitted to The Wistar Institute Genomics Facility for initial analysis of RNA quality, with each sample having a RIN value greater than 8.5 (TapeStation, Agilent Technologies). Sequencing library preparation was then completed using the QuantSeq 3’-mRNA-Seq Library Prep kit (Lexogen) to generate Illumina-compatible sequencing libraries according to the manufacturer’s instructions. Single reads of 75 bp were obtained using a NextSeq 500 sequencer. RNA-seq data were aligned against the Hg19 version of the human genome using Bowtie 2 (67) and RSEM v1.2.12 software (68) was used to estimate raw read counts. DESeq2 (69) was used to estimate significance of differential expression between groups pairs. Genes that passed nominal p<0.05 (FDR<5%) threshold were reported unless stated otherwise. RNA-seq data were deposited for public access at Gene Expression Omnibus. See Data Availability section for accession number.

### Chromatin immunoprecipitation sequencing (ChIP-seq)

Chromatin immunoprecipitation with next-generation sequencing (ChIP-seq) was performed as previously described (31). Briefly, 25 × 10^6^ WT and lamin A/C KO cells per immunoprecipitation were collected and fixed with 1% formaldehyde for 15 min and then quenched with 0.25 M glycine for 5 min on ice. After 3 washes with PBS, pellets were sequentially resuspended in 10 mL each of a series of 2 lysis buffers and in 1mL of the third lysis buffer before fragmentation in Covaris ME220 Ultrasonicator (peak power 75, duty factor 25, cycles/burst 1000, average power 18.8, time 720 s) to generate chromatin fragments roughly 200–500 bp in size. Chromatin was centrifuged to clear debris, and a 1:20 dilution of the cleared chromatin was stored to be used as the standard input for comparison against immunoprecipitations. Chromatin was incubated rotating at 4°C for 1 h with 5 μg H3K9me2 (Abcam, ab1220) and CTCF (Abcam, ab128873) antibodies, then chromatin–antibody complexes were precipitated using 50 μL of Dynabeads Protein A (ThermoFisher, 10001D) incubated rotating at 4°C overnight. DNA was purified using Wizard SV Gel and PCR Clean-up Kit (Promega, A9285). Libraries for sequencing were prepared using NEBNext Ultra II DNA Library Prep Kit (New England Biolabs, E7103) and sequenced on Illumina HiSeq 2500.

### ChIP-seq analysis

Reads were mapped against the human genome assembly using Burrows-Wheeler Aligner (BWA) (70). We used MACS2 (71),(72), (73) software packages to call peaks using input samples as control. deepTools (74) was used for data visualization. For comparison, alignment files were read-depth normalized, and replicates were merged for visualization purposes. Chip-seq data were deposited for public access at Gene Expression Omnibus. See Data Availability section for the accession number.

### Chromatin immunoprecipitation (ChIP)

Chromatin immunoprecipitation was performed as previously described (31) with minor changes. Briefly, 1 × 10^6^ WT and lamin A/C KO cells per immunoprecipitation were collected and fixed with 1% formaldehyde for 15 min and then quenched with 0.125 M glycine for 5 min on ice. After centrifugation, the pellet was resuspended in 120 μL lysis buffer and sonicated using Covaris ME220 Ultrasonicator (peak power 75, duty factor 25, cycles/burst 1000, average power 18.8, time 600 s) to generate chromatin fragments roughly 100–200 bp in size. A sample of ‘input chromatin’ was collected at this point as a standard for comparison against immunoprecipitations (5% of total material). Chromatin was then incubated overnight, rotating at 4°C with 4 μg of antibody against CTCF (Abcam, ab128873), 5 μg of antibody against H3K9me2 (Abcam, ab1220), or 5 μg of IgG antibody (Cell Signaling, 2729). Real-time PCR was performed with a master mix containing 1X Maxima SYBR Green (ThermoFisher Scientific, K0223), 0.25 μM primers, and 1:50 of ChIP or input DNA per well. Quantitative PCRs were carried out in triplicate using the ABI StepOne Plus PCR system. Data were analyzed by the ΔΔCT method (where CT is threshold cycle) relative to DNA input. Primer sequences were as follows: G6PD Rev: 5′-GGGAAGCAGAGCGGAAAG-3′; G6PD Fw: 5′-TGCTCCCACCACTCTATGA-3′; SLC25A1 Rev: 5′-GAGCACCTTTGCCTGGATAA-3′; SLC25A1 Fw: 5′-CAGGCCAGTTTGAAGACTACAT-3′; GAD1 Rev: 5′-CAGGAAAGCCCTCATTGTCT-3′; GAD1 Fw: 5′-GAAGTAAGGGCGACGTGAAA-3′; PFKL Rev: 5′-TCCTTCCTCAGTCCTGGAA-3′; PFKL Fw: 5′-TTTACAGCCGAGCACTGAC-3′; ZNF385A Rev: 5′-CGTGGAATCTCGAAACACATTAC-3′; ZNF385A Fw: 5′-AATTTGCCACAACTCCACAAG-3′; HIC1 Rev: 5′-AGGGCGTCTTTGCTGTG-3′; HIC1 Fw: 5′-CAGGAAGTGCGTGGTCTG-3′

### HiC assay (Chromatin conformation capture assay)

Hi-C assay was performed as previously described (31). Briefly, 5 × 10^6^ cells per condition were collected for *in-situ* Hi-C. Libraries of total ligation products were produced using Ultralow Library Systems V2 (Tecan Genomics, 0344NB-32) as per the manufacturer’s protocol. Libraries were sequenced using the Illumina HiSeq 2500 sequencing platform with paired-end 75 bp read length. HiC data were preprocessed using HiC-Pro v2.10.0 pipeline (75) with default settings using the human genome at 1 kb resolution. DESeq2 (69) was used to estimate the significance of differential contact based on raw count matrix files. To identify cell-type-specific loops, we summed all the interactions that passed the FDR threshold based on cell type and further filtered them by CTCF binding. Then a differential analysis was performed using the DESeq2 R package as described above. The detailed protocol with all minor alterations will be supplied by the corresponding author upon request. Hi-seq data were deposited for public access at Gene Expression Omnibus. See Data Availability section for accession number.

### Metabolomic analysis

Metabolomics analyses were performed at The Wistar Institute Proteomics and Metabolomics Shared Resource. WT and lamin A/C KO cells were cultured in glucose-free RPMI-1640 media supplemented with 11.1 mM uniformly labeled U-13C-glucose (Cambridge Isotope Laboratories Inc., D-Glucose-U13C6, 99%, CLM-1396) for 24 hours to reach isotopic steady state. Unlabeled controls were included for each condition. Polar metabolites were extracted using ice-cold extraction solution containing 80:20 (v/v) MeOH/water. Samples were analyzed by liquid chromatography-tandem mass spectrometry (LC-MS/MS) on a Thermo Scientific Q Exactive HF-X mass spectrometer with a HESI II probe in-line with a Vanquish UHPLC system. Samples were analyzed in a pseudorandomized order. LC separation was performed under HILIC conditions using a ZIC-pHILIC column (150 × 2.1 mm, 5 μM) maintained at 45°C (EMD Millipore). Mobile phase A was 20 mM ammonium carbonate, 0.1% ammonium hydroxide, pH 9.2, and mobile phase B was acetonitrile. Analytical separation was performed at a 0.2 mL/min flow rate using the following gradient: 0 min, 85% B; 2 min, 85% B; 17 min, 20% B; 17.1 min, 85% B; and 26 min, 85% B. Samples were analyzed by either full MS scans with polarity switching (all samples) or full MS with data-dependent MS/MS scans, with separate acquisitions for positive and negative polarities (pool of unlabeled samples). Relevant MS parameters included: sheath gas, 40; auxiliary gas, 10; sweep gas, 2; auxiliary gas heater temperature, 350°C; spray voltage, 3.5/3.2 kV for positive/negative polarities; capillary temperature, 325°C; S-lens RF, 40. Full MS scans were acquired with a scan range of 65 to 975 m/z; 120,000 resolution; automated gain control (AGC) target of 1E6; and a maximum injection time (IT) of 100 ms. Data-dependent MS/MS was performed on the 10 most abundant ions; 15,000 resolution; AGC target of 5E4; maximum IT of 50 ms; isolation width of 1.0 m/z; and stepped normalized collision energy of 20, 40, and 60. Raw data were processed using Compound Discoverer 3.1 (Thermo Scientific) with separate analyses for positive and negative polarities. Metabolites were identified by matching accurate mass and retention time to standards or by querying MS/MS scans against the mzCloud spectral database (full match, score > 50; mzCloud.org). Identifications were transferred to labeled samples in isotope tracing experiments, considering all possible 13C isotopologues. Metabolite levels were quantified by integrated peak area for all detected adducts using full MS only data. Isotope tracing data were corrected for natural ^13^C abundance and isotope tracer purity. Metabolite levels were further normalized to total protein recovered from the polar metabolite extraction pellet.

### Droplet digital PCR (ddPCR)

DNA was isolated from 1×10^6^ cells using the GeneJET Genomic DNA Purification Kit (ThermoFisher Scientific, K0722) according to the manufacturer’s protocol. 500 ng of DNA was digested with BamHI enzyme (10 U/μl, New England Biolabs) in a total volume of 10 μL for 1 h at 37°C. The digestion reaction was then diluted 1:20 in nuclease-free water. 10 μL of diluted DNA digest was mixed with 12.5 μL of 2X digital PCR supermix for probes (no dUTP) (Bio-Rad), 1.25 μL 20X FAM primers, and 1.25 μL 20X VIC primers for each reaction. Primer and probe sequences for EBV Lmp1 were as follows: Fw (5′-3′) AAGGTCAAAGAACAAGGCCAAG, Rv (5′-3′) GCATCGGAGTCGGTGG, and probe FAM-AGCGTGTCCCCGTGGAGG. Host control primer and probe sequences for ribonuclease P protein subunit 30 (Rpp30) were as follows: Fw (5′-3′) GATTTGGACCTGCGAGCG, Rv (5′-3′) GCGGCTGTCTCCACAAGT, and probe VIC-CTGACCTGAAGGCTCT. 20X primers contained 18 μM PCR primers and 5 μM probes for a final PCR reaction concentration of 900 nM PCR primers and 250 nM probe. Each sample was run in duplicate. The ddPCR plate was sealed with a foil heat seal using the PX1 PCR Plate Sealer (Bio-Rad) at 180°C for 5 s. The plate was vortexed and spun down at 1,000 rpm for 1 min. Droplets were generated using the QX200 Droplet Digital PCR System (Bio-Rad), and transfer of emulsified samples to a PCR plate was performed according to the manufacturer’s instructions. A PCR plate containing emulsified droplets was sealed with a foil heat seal. PCR reactions were performed on the C1000 Touch Thermal Cycler (Bio-Rad). The cycling protocol included an enzyme activation step at 95°C for 10 min and cycled 40 times between a denaturing step at 94°C for 30 s and an annealing and extension step at 60°C for 1 min. Finally, one enzyme deactivation step was performed at 98°C for 10 min. The ramp rate between these steps was set at 2°C/s. Droplets were then counted using the QX200 Droplet Reader (Bio-Rad). The absolute quantity of DNA per sample was determined using the QuantaSoft software.

### Proximity ligation assay (PLA)

Proximity ligation assay (PLA) was performed using the Duolink PLA Fluorescence kit (Sigma Aldrich) according to the manufacturer’s protocol. CTCF, lamin B1, and H3K9me2 antibodies were used, and images were acquired at The Wistar Institute Imaging core.

### Histone extraction

Cells were harvested and washed twice using ice-cold PBS. Cells were resuspended at a cell density of 10^7^ cells per mL in Triton extraction buffer (TEB) consisting of PBS containing 0.5% Triton X-100 (v/v), 2 mM phenylmethyl sulfonyl fluoride (PMSF), 0.02% (w/v) sodium azide (NaN3) Cells were lysed on ice for 10 minutes with gentle stirring, centrifuged at 2,000 rpm for 10 minutes at 4°C. The supernatant was removed and discarded. Cells were washed in half the volume of TEB and centrifuged at the same speed as before. Pellets were resuspended in 0.2 N HCl at a cell density of 4 × 10^7^ cells per mL. Acid extraction of histones was performed overnight at 4 °C. The next day, samples were centrifuged at 2,000 rpm for 10 minutes at 4 °C. The supernatant was moved to a clean tube, and the protein content was determined by Bradford assay.

### Drug screening

Test compounds (1,000X concentration in 100% DMSO) were added in 25 nL aliquots using the Echo 650 acoustic liquid handler to white, opaque tissue culture-treated 384-well plates. Cells were seeded in 25 μL of complete media (RPMI + 10% FBS + pen/strep) at 1,000 cells/well, resulting in a final DMSO concentration of 0.1%. Each compound was tested at 10, 1, 0.1, and 0.01 μM. Bortezomib treatment (1 μM) was included as positive control. After 5 days of incubation at 37°C + 5% CO_2_, cell viability was assessed by adding 10 μL of CellTiter-Glo (Promega, G7570) and measuring luminescence. The relative light unit (RLU) values were converted to % toxicity, with 100% corresponding to the RLU value in the presence of bortezomib and 0% to the RLU value of control samples (0.1% DMSO). The % toxicity values at each concentration of compound were fit to a 4-parameter dose-response equation using XLFit5 (IDBS) with the top and bottom values fixed to 100% and 0%, respectively.

### Cell viability

Cell viability was measured using the LIVE/DEAD Viability/Cytotoxicity Kit (ThermoFisher Scientific, L3224). Cell staining was performed according to the manufacturer’s protocol. Images were acquired using the EVOS M5000 Microscope. RFP, GFP, and brightfield images were acquired for each sample in triplicate. Cell counting was performed with ImageJ software using the Analyze Particle option.

### Seahorse XF Cell Mito Stress Test assay

Seahorse XF cell plates (XF96, Agilent 103794-100) were coated with Cell-Tak Cell and Tissue Adhesive solution (Corning, 354240) at a concentration of 22.4 μg/mL (diluted in 0.1 M sodium bicarbonate, pH8.0) to allow for adhesion of suspension cells to the plates. Cells were then seeded (2 × 10^5^/well) in the XF96 plates, followed by centrifugation at room temperature at 200 *×* g for 5 minutes. The plated cells were then incubated in a 37°C incubator not supplemented with CO_2_ for 25–30 minutes to ensure that the cells had completely attached. Cells were incubated for a total of 1 hr in a 37°C incubator without CO_2_ to allow for pre-equilibration with the assay medium. Cells were then analyzed by the XF Cell Mito Stress Test assay to measure the oxygen consumption rate (OCR) and extracellular acidification rate (ECAR) of WT and lamin A/C KO cells in XF media (non-buffered RPMI containing 10 mM glucose, 2 mM L-glutamine, and 1 mM sodium pyruvate). OCR and ECAR were detected under basal conditions followed by the sequential addition of 2 μM oligomycin (Sigma), 1 μM fluoro-carbonyl cyanide phenylhydrazone (FCCP, Sigma), and 2 μM rotenone + 2 μM antimycin A (Sigma). This allowed for an estimation of the contribution of individual parameters to basal respiration, proton leak, maximal respiration, spare respiratory capacity, non-mitochondrial respiration, and ATP production. The XF Mito Stress Test report generator automatically calculated the parameters from Wave software (Agilent) data that were exported to Excel. All experiments were performed with 4 to 10 replicate wells in the XF96 Extracellular Flux Analyzer (Agilent).

### Statistical analysis

All experiments presented were conducted at least in triplicate to ensure the reproducibility of results. The Prism (GraphPad) statistical software package was used to identify statistically significant differences between experimental conditions and control samples, using Student’s *t*-test.

## RESULTS

### Lamin A/C expression impacts gene expression, phenotype, and metabolism of B cells

Our previous research demonstrated that EBV infection in B cells induces the expression of lamin A/C, which is critical in regulating viral latent gene expression (56). We hypothesized that lamin A/C increased expression may also impact global gene expression in infected B cells. To test our hypothesis, we used RNA-Seq to analyze the global transcriptome of EBV+ LCL WT cells (hereafter WT) and EBV+ LCL cells in which expression of lamin A/C was knocked out via CRISPR/Cas9 (hereafter lamin A/C KO or KO cells). These cells show no differences compared to WT cells in terms of lamin B1 expression or the number of EBV genomes per cell (**Fig. S1A and S1B**). Principal component analysis (PCA) revealed a clear separation of the datasets depending on the presence of lamin A/C, with principal component 1 (PC1, corresponding to lamin A/C deletion) accounting for 67.04% of the observed variance between conditions (**Fig. 1A**). By differential expression gene analysis, we identified 5,643 genes that were deregulated in lamin A/C KO cells compared with WT cells, of which 4,185 genes (74%) were upregulated, and 1,458 (26%) were downregulated (**Fig. 1B**). These data are consistent with the role of lamin A/C in regulating transcription (76). Among the most upregulated transcripts in lamin A/C KO cells, we observed several pseudogenes and long non-coding RNAs (lncRNAs), including metastasis-associated lung adenocarcinoma transcript 1 (MALAT1, **Fig. 1C**), a well-studied lncRNA associated with immune evasion in diffuse large B-cell lymphoma (77). The most downregulated genes in KO cells included the leukocyte tyrosine kinase (LTK) gene (**Fig. 1C**), which is expressed in lymphocytes and monocytes and is involved in the development of B lymphocytes (78). These observations suggest that lamin A/C expression may play a role in the activation and differentiation of B cells. Furthermore, by employing the CIBERSORT software (79,80) to computationally infer the proportions of distinct immune cell types between WT and lamin A/C KO cells based on their transcriptomic profiles, we found that WT cells predominantly displayed the phenotype of memory B cells, while KO cells exhibited characteristics akin to naïve B cells (**Fig. 1D**), further indicating that lamin A/C expression likely influences B-cell differentiation. In support of this observation, we assessed lamin A/C expression in memory B cells and naive B cells using expression data from the Human Protein Atlas (HPA) tissue sample RNA-Seq dataset (81) and observed that lamin A/C was expressed in memory B cells but not in naïve B cells (**Fig. 1E**). To gain a deeper insight into the biological significance of lamin A/C-mediated gene regulation, we used Ingenuity Pathway Analysis (IPA) (82) to identify the cellular pathways and regulators affected by lamin A/C expression (**Fig. S2**). The IPA analysis indicated that lamin A/C expression exerts influence on multiple cellular pathways pertinent to B-cell biology (**Fig. S2A**). Among the pathways upregulated in lamin A/C KO cells, we identified the PI3K signaling pathway, which plays a critical role in B-cell development, activation, differentiation, and generation of memory B cells (83,84); the ARP2/3-WASP complex, which has implications for B-cell spreading (85,86); and the unfolded protein response (UPR), which is essential for plasma B cells (**Fig. S2A**) (87,88). The absence of lamin A/C resulted in suppression of 2 essential metabolic pathways—glycolysis and gluconeogenesis —both of which play a critical role in cell proliferation and differentiation (**Fig. S2A**) (89). The IPA analysis also uncovered several potential upstream transcriptional regulators that might be involved in driving the observed changes in gene expression within the KO cells. Specifically, the analysis suggested that transcription factors important for B-cell activation, proliferation, and survival, such as IRF4, IRF7, and REL(90–93), and transcription factors pivotal in cell metabolism, such as ATF4 and TCF7L2 (94,95), may be activated in lamin A/C KO cells (**Fig. S2B**). Conversely, upstream regulator analysis predicted inhibition of NUPR1, HIF1A, PHF12, and KLF3 (96–99), indicating reduced stress tolerance, impaired metabolic adaptation, and weakened chromatin repression. Together, these changes suggest that loss of lamin A/C compromises key transcriptional networks required for B-cell activation, survival, and maintenance of lineage-appropriate gene expression (**Fig. S2B**). Overall, our transcriptomic analysis indicates that lamin A/C KO B cells undergo a coordinated transcriptional rewiring that reduces their ability to maintain homeostasis while amplifying stress-and activation-driven pathways.

**Figure 1.**
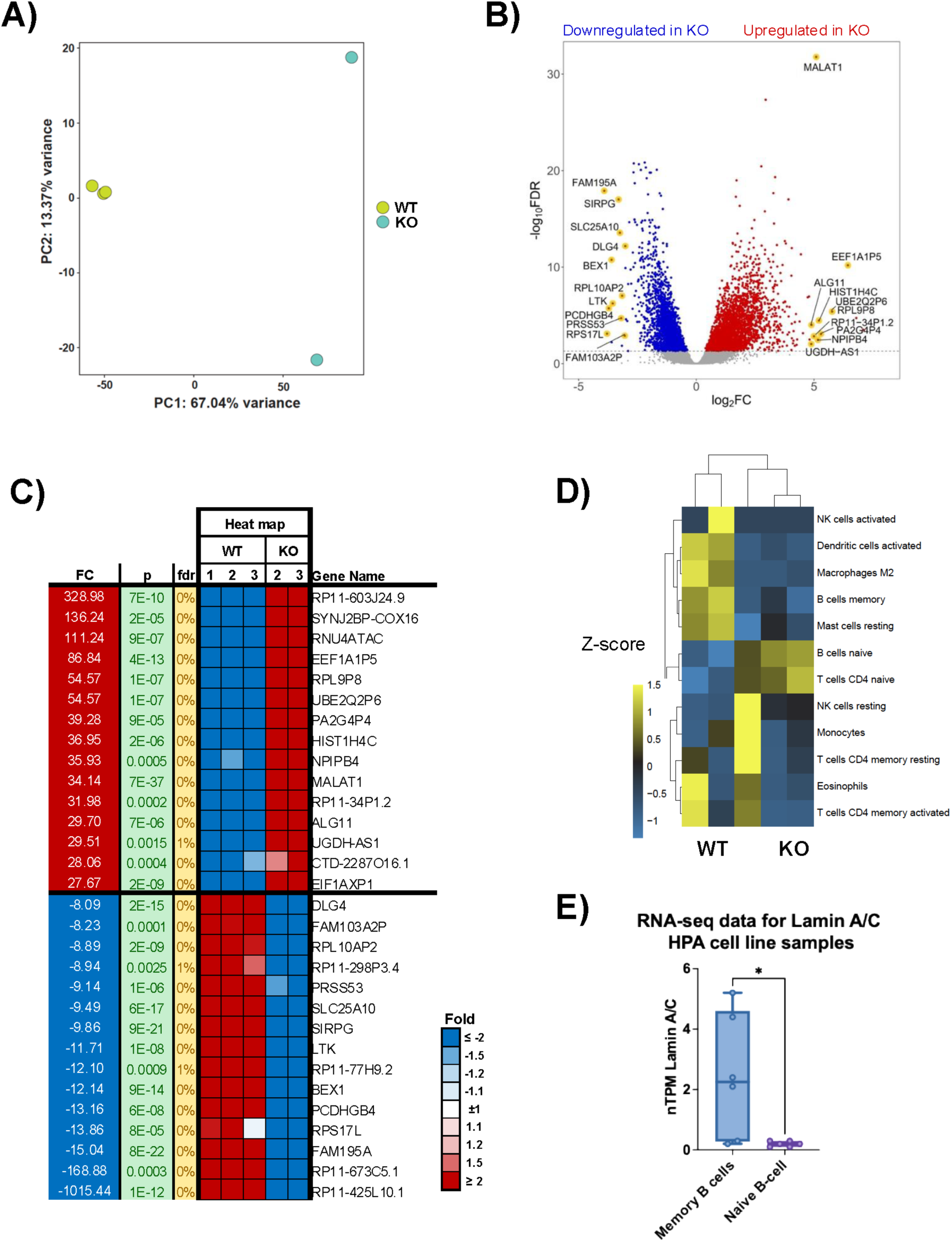
**Lamin A/C knockout alters host gene expression in EBV+ LCL cells**. **A)** Principal component analysis (PCA) scatter plot of RNA-Seq data from lamin A/C KO and WT LCL cells. The samples are shown as a function of principal component 1 (PC1, x-axis) and principal component 2 (PC2, y-axis). The percentage of variance accounted for by PC1 and PC2 is indicated. **B)** Differentially expressed genes (false discovery rate [FDR] < 5%, |log2 FC| > 1.5) in lamin A/C KO cells compared with WT cells. Upregulated genes are depicted in red, while downregulated genes are depicted in blue. Genes with the highest change in expression are highlighted in yellow. **C)** Heat map of RNA-Seq data from WT and lamin A/C KO cells showing the 30 top regulated genes with higher and lower fold change (FC) and FDR <5%, p<0.05. **D)** Relative percentage of different immune cell type profiles in WT and lamin A/C KO cells. The immune cell profiles were deconvoluted from the RNA-Seq data using CIBERSORT. **E)** Lamin A/C abundance in memory and naive B cells based on expression data from the Human Protein Atlas (HPA) tissue sample RNA-Seq dataset. nTPM, normalized transcripts per million; *, p < 0,01.

### Lamin A/C depletion causes reorganization of the chromatin architecture, affecting gene expression

The interaction between the nuclear lamina and chromatin regions is involved in chromatin looping (100,101). To investigate the mechanism by which lamin A/C regulates gene expression, we focused our attention on chromatin structure and position within the nuclear space and hypothesized that depletion of lamin A/C may affect the 3D structure of the genome. We used Hi-C assay to generate maps of the chromatin loop interactions in WT and KO cells. By comparing these maps, we observed that the depletion of lamin A/C had a significant effect on global genome organization, with 39 compartments switching from B (inactive chromatin) to A (active chromatin) compartments and 23 from A to B in lamin A/C KO cells compared with the WT sample (**Fig. 2A**). Next, we analyzed the Hi-C datasets to assess changes in genome organization occurring at the level of chromatin loops in KO cells. We observed an increased frequency of smaller chromatin loops (occurring between nearby regions) and a decreased occurrence of larger loops (connecting distant regions) in KO cells compared with WT cells (**Fig. 2B**). To assess whether changes in higher-order chromatin organization were associated with transcriptional alterations in Lamin A/C KO cells, we integrated RNA-seq and Hi-C datasets. Transcriptionally deregulated genes were then overlapped with genomic regions exhibiting altered chromatin looping as determined by Hi-C. We identified 270 genes located within the modified chromatin loops whose expression was altered in KO cells compared with WT (**Fig. 2C**). Notably, our analysis uncovered a consistent pattern: the loss of a chromatin loop in KO cells was associated with a decrease in gene transcription (**Fig. 2D**), while the acquisition of a chromatin loop consistently led to transcriptional activation (**Fig. 2E**), highlighting the dynamic interplay between lamin A/C, chromatin organization, and gene expression. Together, these data indicate that KO-induced remodeling of chromatin looping is closely linked to changes in gene expression

**Figure 2.**
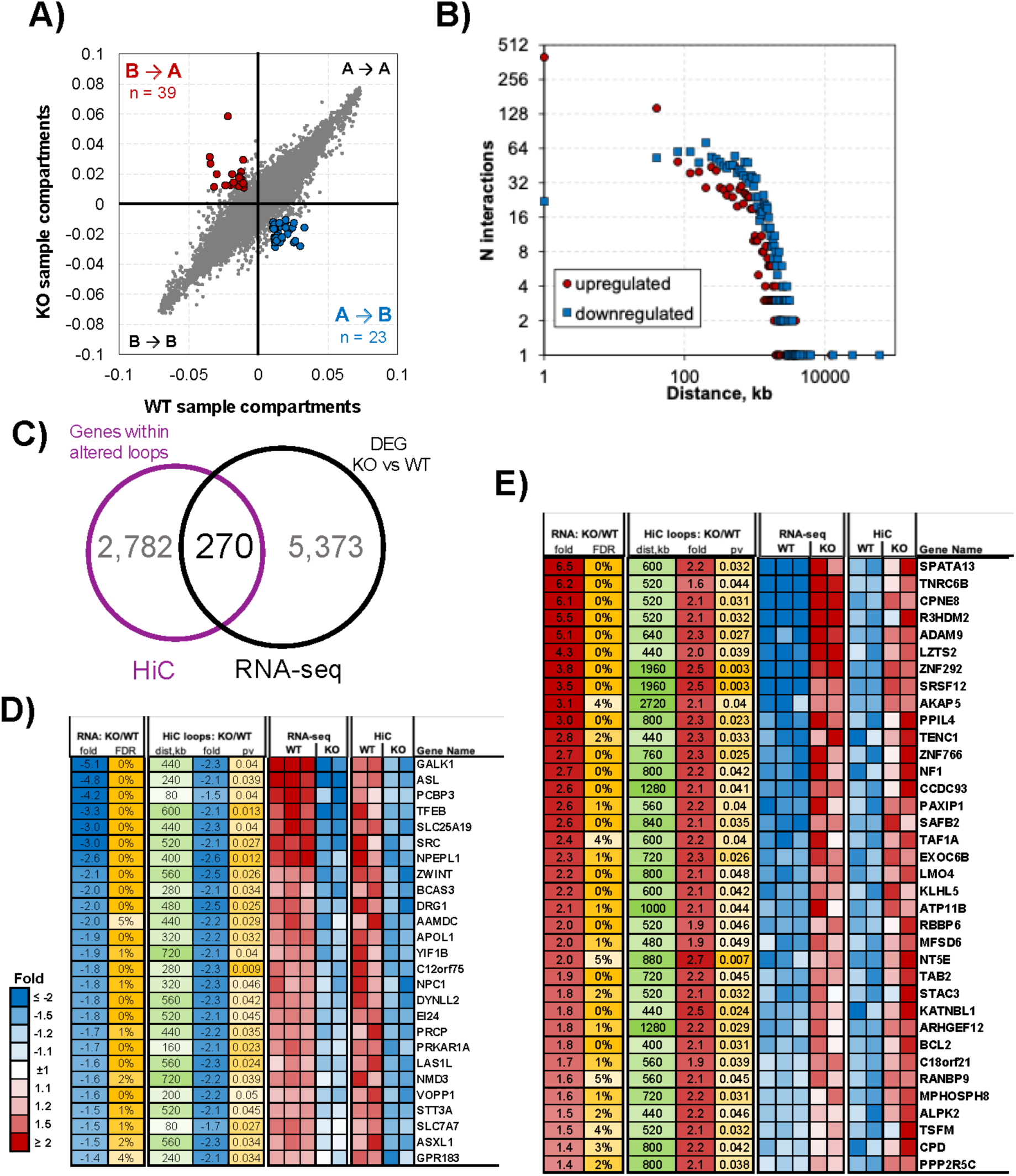
Depletion of lamin A/C affects the genome architecture of EBV+ B cells. **A)** Eigenvector in WT versus lamin A/C KO cells from Hi-C data sets showing significant (p < 0.01) changes in open *(A)* and closed chromatin (*B)* compartments (compartment switch). Significantly changed compartments are depicted in red (*B* to *A*, n = 39) and blue (*A* to *B*, n = 23). **B)** Number of significant (p < 0.05) Hi-C interactions represented as a function of distance (measured in kb) in lamin A/C KO cells compared with WT cells. **C)** Graph showing the overlap between the group of genes located in proximity of altered chromatin loops (Hi-C dataset) and those whose transcription was affected by lamin A/C ablation (RNA-Seq dataset). DEG, deregulated genes. **D)** and **E)** Heat maps of changes in gene expression and chromatin looping in downregulated (D) and upregulated (E) genes in KO cells compared with WT cells. Genes with a false discovery rate (FDR) < 5% and p < 0.05 are shown, and fold change values are represented in the table.

### Lamin A/C-deregulated genes are bound by CTCF

Previous studies have shown that CTCF occupancy on the human genome occurs at many lamina-associated domains (LADs) (102,103), indicating a potential interaction between CTCF and nuclear lamin proteins in the regulation of gene expression. To explore if lamin A/C depletion was associated with changes in CTCF occupancy across the host genome, we used ChIP-Seq to compare binding in WT cells and lamin A/C KO cells and observed no alterations in either the peak intensity or localization of CTCF (**Fig. 3A**). We next analyzed CTCF binding among the genes deregulated by lamin A/C depletion by cross-comparing CTCF ChIP-Seq data from WT cells with our RNA-Seq datasets and identified 328 genes common to both datasets, representing a significant 1.23-fold enrichment over random expectation (**Fig. 3B)**. This significant overrepresentation of CTCF-bound genes among the genes downregulated in lamin A/C KO cells may suggest an interplay between CTCF and lamin A/C in regulating gene expression. IPA analysis of these common lamin A/C-CTCF targets showed that inactivated pathways included glycolysis, the AMP-activated protein kinase (AMPK) signaling, and sirtuin signaling, while activated pathways included the mitochondrial dysfunction pathway and granzyme A signaling (104), which is linked to B-cell differentiation (**Fig. 3C**). Additionally, we identified several transcription factors whose targets were overrepresented among the lamin A/C-CTCF deregulated genes, such as PAX5 and KLF3 (activated) (105,106), both of which play a critical role in B-cell differentiation, and MYC and POU2F2 (inhibited)(107–110), which are critical for controlling the proliferation and differentiation of B cells and are typically deregulated in lymphomas and EBV-driven malignancies (**Fig. 3D**). Overall, these results suggest that the effect of lamin A/C on gene expression and regulation of B-cell biology may be mediated, at least in part, by CTCF functions.

**Figure 3:**
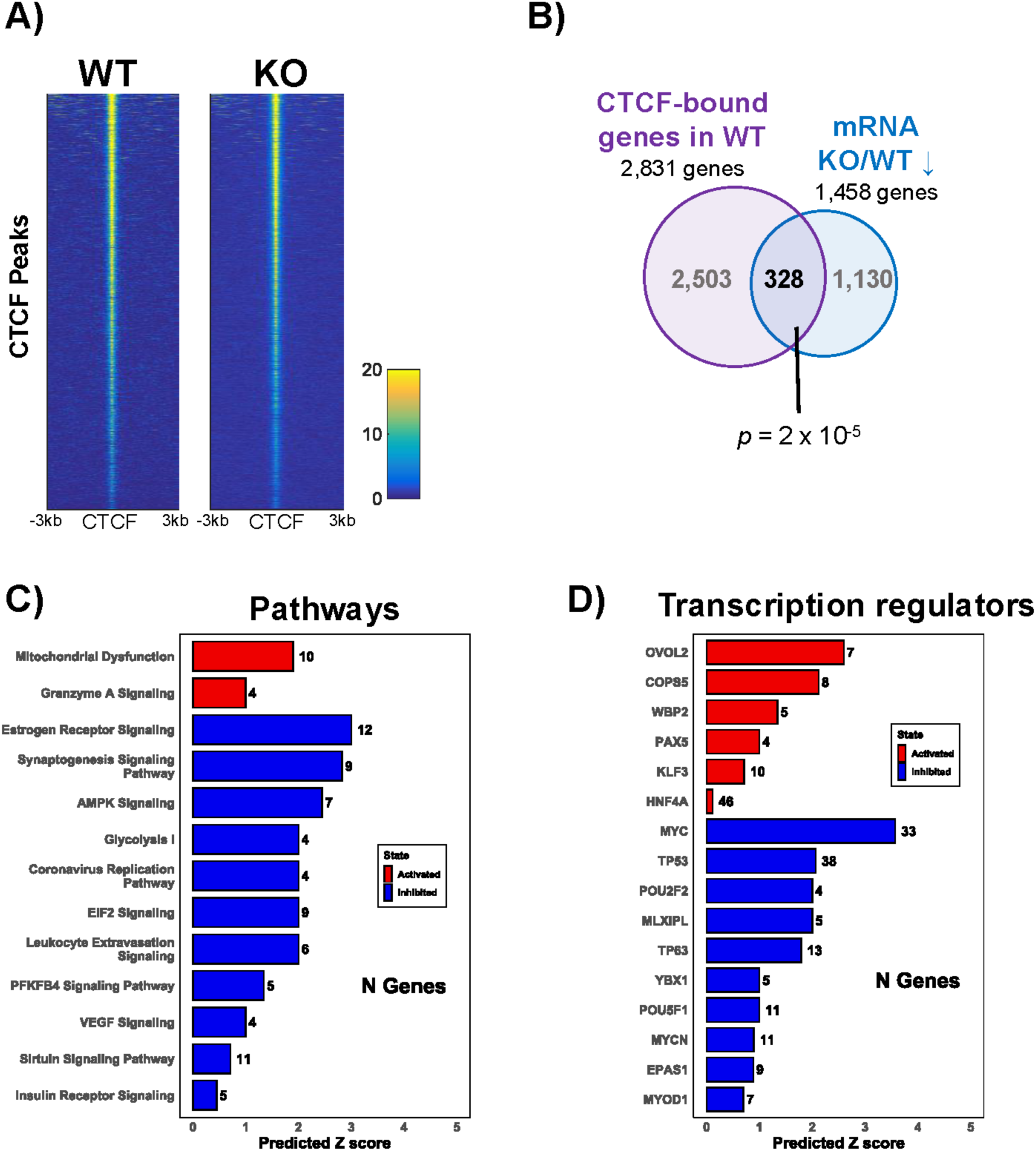
CTCF occupancy on genes deregulated by lamin A/C KO. **A)** CTCF ChIP-Seq peaks ± 3 kb in WT and lamin A/C KO cells. Each row represents a single peak; the intensity of the signal was measured as RPKM, a normalized number of reads. Yellow and blue represent high-and low-read densities, respectively. **B)** Overlapping CTCF-bound genes (identified by ChIP-Seq) and genes downregulated in lamin A/C KO cells compared to WT (identified by RNA-Seq). The overlap between the two conditions is statistically significant (p = 2 × 10^-5^). **C** and **D)** Ingenuity Pathway Analysis (IPA) of the 328 overlapping genes showing the top deregulated pathways (C) and upstream transcriptional regulators (D).

### CTCF relocalization at the nuclear periphery mediates the role of lamin A/C in gene regulation

Since there was no change in global CTCF binding in lamin A/C KO cells, we hypothesized that depletion of lamin A/C may affect CTCF function by impacting its localization within the nuclear space. We used the proximity ligation assay (PLA) in WT and lamin A/C KO cells to assess the interaction between CTCF and 2 well-established markers for nuclear periphery: lamin B1 and H3K9me2 (48,111), whose levels are not affected by lamin A/C depletion in B cells **(Figs. S1A**). We observed increased interaction of CTCF with both lamin B1 and H3K9me2 in the absence of lamin A/C (**Fig. 4A**). This observation suggests that in absence of lamin A/C, CTCF became repositioned towards the nuclear periphery, a location typically associated with gene repression and heterochromatin formation. We therefore hypothesized that the repression of CTCF target genes in lamin A/C KO cells may result from increased deposition of heterochromatin marks, specifically the nuclear periphery–associated modification H3K9me2. To study this, we used ChIP-Seq assay to assess H3K9me2 deposition at CTCF-bound targets deregulated by lamin A/C depletion. We observed an increase in H3K9me2 deposition upstream of the transcription start site (TSS) and across the gene body of the CTCF-bound genes in KO cells compared with WT cells (**Fig. 4B**), demonstrating a strong correlation between H3K9me2 enrichment and decrease in gene transcription. To confirm the ChIP-Seq analysis, we used quantitative ChIP to assess H3K9me2 incorporation at the promoter region of a subset of CTCF-bound deregulated genes. Consistent with ChIP-seq data, we found a significant enrichment of H3K9me2 deposition in KO cells compared with WT cells (**Fig. 4C**). Together, these findings indicate that lamin A/C plays a role in regulating gene expression by affecting CTCF nuclear localization and H3K9me2 deposition at regulated genomic regions.

**Figure 4:**
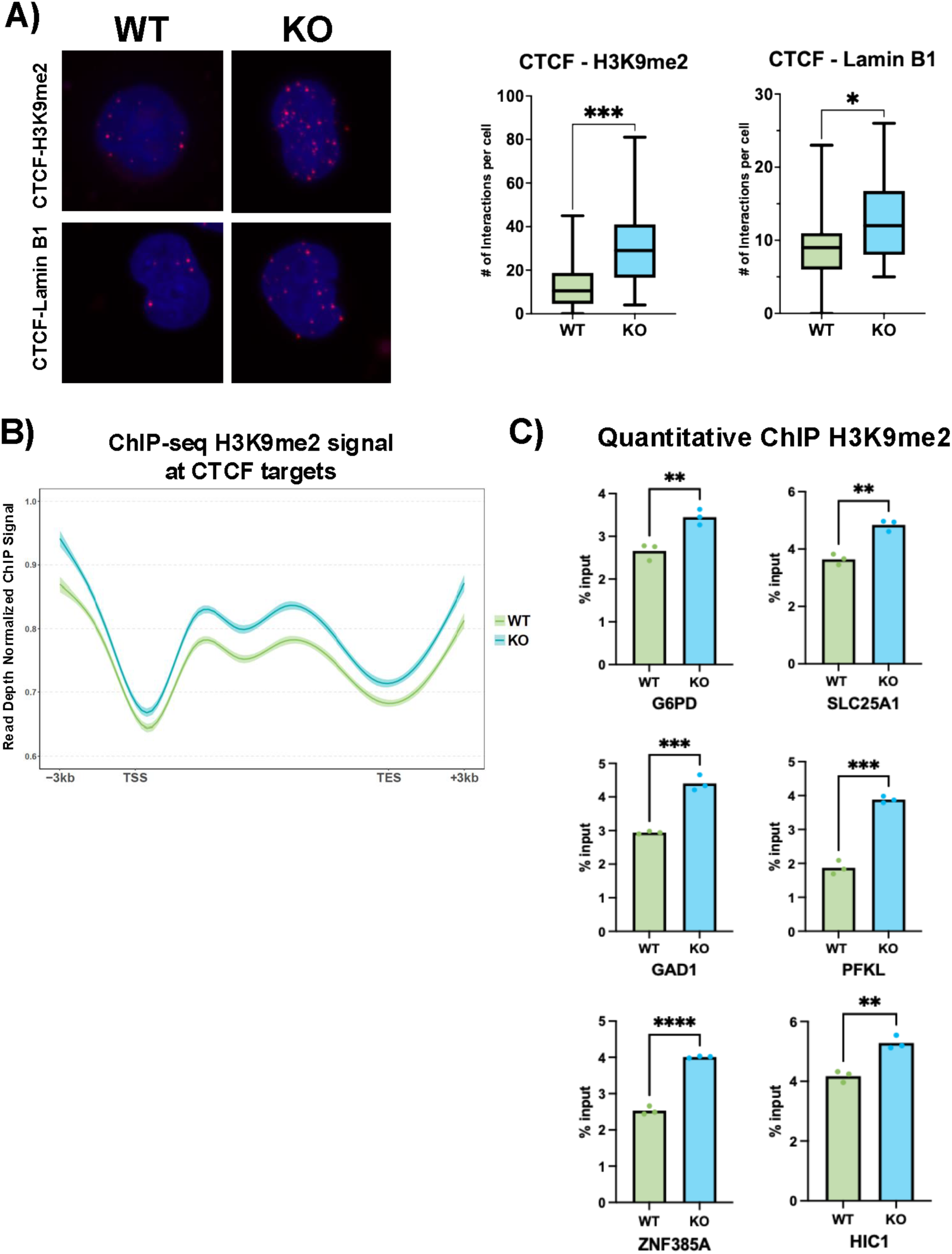
Depletion of lamin A/C causes repositioning of CTCF at the nuclear periphery. **A)** Representative confocal images of in situ proximity ligation assay (PLA) assessing the interaction of CTCF with di-methyl histone H3 lysin 9 (H3K9me2, top panels) and lamin B1 (bottom panels) in WT (left panels) and lamin A/C KO (right panels) cell lines. Maximum intensity projections of a confocal z-stack, including a whole cell, were performed to observe the maximum amount of PLA puncta (red) and the nuclear region stained by DAPI (blue); magnification: 63x. The graph on the right shows the number of PLA puncta per cell in the nuclear region calculated from 3 different images. Median, minimum, and maximum values are shown. *, p < 0,01 and ***, p < 0,0001. **B)** H3K9me2 ChIP-Seq signal at CTCF targets for WT (green) and lamin A/C KO (blue) cells. Peaks at ± 3 kb from the transcription starting site (TSS) and transcription end site (TES). **C)** Quantitative ChIP analysis in WT and lamin A/C KO cells, showing H3K9me2 deposition at the promoter regions of selected CTCF-bound genes deregulated in KO cells. Values are expressed as % of the input. **, p ≤ 0.01; ***, p ≤ 0.001; ****, p ≤ 0.0001.

### Inhibition of H3K9 dimethylation phenocopies lamin A/C depletion

Our data thus far suggested that the changes in gene expression associated with lamin A/C ablation could be due to a repositioning of chromatin regions within the nucleus. Given that contacts between chromatin and the nuclear lamina are promoted by the H3K9 methyltransferase G9a through H3K9me2 deposition (112–114), we tested whether inhibition of G9a may mimic the effect of lamin A/C knockdown on gene expression by treating WT cells with UNC0631, a potent and specific G9a inhibitor (G9ai) (PMC11980580) to lower the global level of H3K9me2 (**Fig. S3A**). We then analyzed by RNA-Seq the effects of G9a inhibition on gene expression. By PCA, the RNA-Seq datasets of the untreated cells (control) were separated from those of the UNC0631-treated (G9ai) cells with a 56.11% variance (**Fig. S3B**). Comparison of the control and G9ai datasets revealed 2,346 deregulated genes (FDR < 5%), **(Fig. S4C).** Among the most upregulated and down-regulated genes were MALAT1 and LTK, respectively, which were also found among the deregulated genes in lamin A/C KO samples (**Fig. S3D**). IPA showed that genes deregulated by G9ai treatment were enriched for specific metabolic and cellular signaling pathways, including the glycolysis and gluconeogenesis pathways among the most inhibited pathways and the PI3K/AKT, ATM, and Jak/STAT signaling pathways among the most activated pathways (**Fig**. **S3E**). Genes involved in glycolysis and gluconeogenesis and the PI3K-AKT signaling pathway were also overrepresented among the deregulated genes in lamin A/C KO cells (**Fig. S2A**). IPA analysis of RNA-Seq datasets for transcription factors showed that target genes of TCF7L2, NUPR1, HIF1A, MYC, and KLF3 were overrepresented among the genes deregulated by G9a inhibition (**Fig. S3F**). Some of these transcription factors were also found concordantly activated or inactivated in lamin A/C KO cells (**Fig. S2B**). These overlaps are suggestive of an interplay between H3K9me2 deposition and lamin A/C in the regulation of gene expression at specific genomic regions. Based on these observations, we predicted that blocking H3K9me2 deposition by G9a inhibition would produce comparable effects to lamin A/C depletion on global gene expression. Therefore, we assessed the overlap between differentially expressed genes upon lamin A/C KO and G9ai treatment and found 1,857 differentially expressed genes in common between the two conditions, corresponding to a 2.5-fold higher overlap than expected by chance (**Fig. 5A**). These common genes represented approximately 80% of all differentially expressed genes inG9a inhibitor-treated cells and 30% of all the differentially expressed genes in lamin A/C KO cells. Additionally, a significant number of genes were concordantly regulated between lamin A/C KO and G9ai treatment (930 genes upregulated in both conditions and 927 genes downregulated in both conditions, **Fig. 5B**), with comparable fold changes (**Fig. S4A** and **S4B**). The top 30 upregulated and downregulated genes are shown in **Fig. S4C** and **S4D**, respectively. Analysis of H3K9me2 deposition across the gene body of the 1,857 common targets in ChIP-Seq datasets of WT and lamin A/C KO cells revealed an increased deposition of H3K9me2 mark across the gene body of the 927 targets that were downregulated in KO cells compared with WT (**Fig. 5C**). In contrast, no significant differences in H3K9me2 deposition were observed across the gene body of the 930 upregulated genes (**Fig. 5D**). IPA of the 1,857 common targets revealed an overrepresentation of genes of the ribonucleotide reductase signaling and the ATM signaling among the activated pathways and an overrepresentation of glycolysis and gluconeogenesis genes among the inhibited pathways (**Fig. 5E**). IPA analysis of transcriptional regulators predicted activation of TCF7L2, VHL, IRF4, and REL and inhibition of HIF1A, KLF3, and STAT4 (**Fig. 5F**). Differential expression analysis of genes that encode proteins involved in PI3K signaling, glycolysis pathway, and MYC-mediated apoptosis pathways showed concordant regulation and similar magnitude of the transcriptional changes driven by G9ai treatment and lamin A/C KO. (**Fig. 5G-I**). Overall, these data indicate that lamin A/C may regulate the expression of genes involved in cellular metabolism and signaling via H3K9me2 deposition.

**Figure 5:**
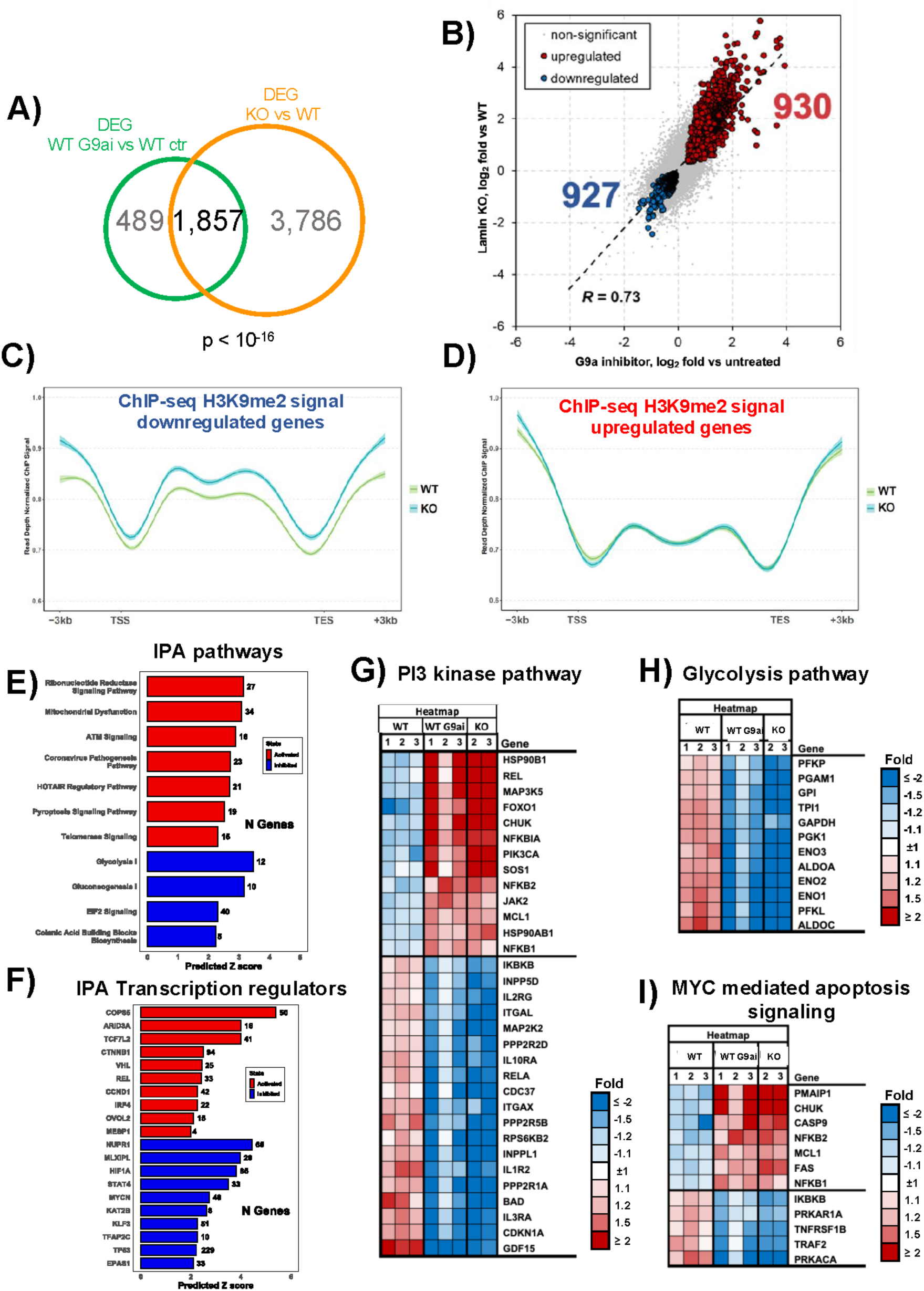
H3K9me2 inhibition affects gene expression in WT cells and phenocopies the effects of lamin A/C KO. **A)** Overlap of deregulated genes (DEG) in RNA-Seq analysis between lamin A/C KO and G9a inhibitor treatment in WT cells. The overlap was statistically significant (p < 10^-16^). **B)** Concordant regulation of the genes whose expression was altered by both lamin A/C KO and G9a inhibition. Data are represented using log_2_ fold changes, and significantly regulated genes are depicted in red (upregulated) and blue (downregulated). The correlation coefficient (R = 0.73) indicates a strong positive correlation between the datasets. **C)** and **D)** H3K9me2 ChIP-Seq signal across the gene body of the 1,857 common targets between lamin A/C KO and G9a inhibition in WT cells. Increased deposition of H3K9me2 mark on the 927 targets that were downregulated in KO cells compared with WT (C), and no significant difference in H3K9me2 deposition on the 930 upregulated genes (D). Peaks at ± 3 kb from the transcription starting site (TSS) and the transcription end site (TES). **E)** and **F)** Ingenuity Pathway Analysis of the 1,857 common targets between lamin A/C KO and G9a inhibitor-treated WT cells, showing the top deregulated pathways (E) and upstream transcriptional regulators (F). Red indicates activation and blue inhibition. Data are shown as a function of the predicted z-score. **G-I)** Heat-maps of RNA-Seq data from WT, G9a inhibitor-treated WT, and lamin A/C KO cells showing genes in the PI3K, glycolysis, and c-myc signaling pathways whose expression was concordantly altered (false discovery rate [FDR] < 5%) by lamin A/C KO and G9a inhibition. Color code represents different fold change (FC) values, with upregulated genes shown in red and downregulated genes shown in blue.

### Silencing of lamin A/C rewires the cellular metabolism of B cells

Our transcriptomic analysis indicates that the rearrangement of chromatin regions within the nucleus, controlled by lamin A/C, may influence the function of transcription factors and the expression of genes involved in cell and cancer metabolism. We hypothesized that lamin A/C depletion may reprogram the cellular metabolism, affecting anabolic and metabolic programs that support aberrant cellular growth. To investigate potential differences in the metabolic requirements of WT and lamin A/C KO cells, we assessed the glycolytic activity and measured the extracellular acidification rate (ECAR) in both conditions. (**Fig. 6A)**. KO cells displayed a lower ECAR level compared with WT cells, indicating a reduced rate of glycolysis. We then evaluated the oxygen consumption rate OCR/ECAR ratio, which serves as an indicator of whether cells favor oxidative phosphorylation (OXPHOS) or glycolysis for energy production. Notably, KO cells exhibited a higher OCR/ECAR ratio compared with WT cells (**Fig. 6A**), suggesting that KO cells rely less on glycolysis (anaerobic metabolism) and instead favour OXPHOS for energy production. This shift away from anaerobic glycolysis suggests a reduced dependence of lamin A/C KO cells on the Warburg effect (115), with pyruvate being channelled to its oxidation into acetyl-CoA for entry into the mitochondrial tricarboxylic acid (TCA) cycle, which provides cancer cells with both energy and biosynthetic precursors essential for their growth and proliferation. We used mass spectrometry metabolic analysis to assess changes in the cellular metabolome between WT and KO cells. Unsupervised PCA of the metabolomic data showed that the absence of lamin A/C in B cells accounted for 82.9% of the variance in metabolomic profile between KO and WT cells (**Fig. S5A**). Lamin A/C silencing induced both upregulation and downregulation of several metabolites (**Fig. 6B, S5B**). Overall, this metabolic analysis suggests that lamin A/C ablation impacts pathways crucial for energy production and synthesis of various biological macromolecules, including glycolysis, purine metabolism, and gluconeogenesis based on metabolite set enrichment analysis (MSEA) conducted using MetaboAnalyst (**Fig. 6C**) (116). Among the pathways most affected by lamin A/C KO, the TCA cycle showed higher abundance of multiple key pathway components in KO cells compared with WT cells (**Fig. 6D, E**). Other pathways that showed a significant enrichment of numerous metabolites in lamin A/C KO cells included the glycine–serine and one-carbon pathway (SGOC), a critical metabolic route supporting nucleotide and lipid synthesis, redox balance, and cell proliferation (**Fig. S6A, B**), and the biosynthesis of several amino acids (**Fig. S6C, D)** (117,118). Interestingly, among the metabolites whose steady-state levels were affected by lamin A/C KO we found many that are involved in the regulation of epigenetic mechanisms, such as the D-2-hydroxyglutarate (D-2HG), which acts by inhibiting key demethylase enzymes (DNA and histone demethylase) (119), or the S-adenosylmethionine (SAM), which is responsible for donating methyl groups to many methyltransferase enzymes responsible for methylating DNA, histones, and RNA (120) and it has been linked to the metabolic changes driven by EBV infection (62).

**Figure 6:**
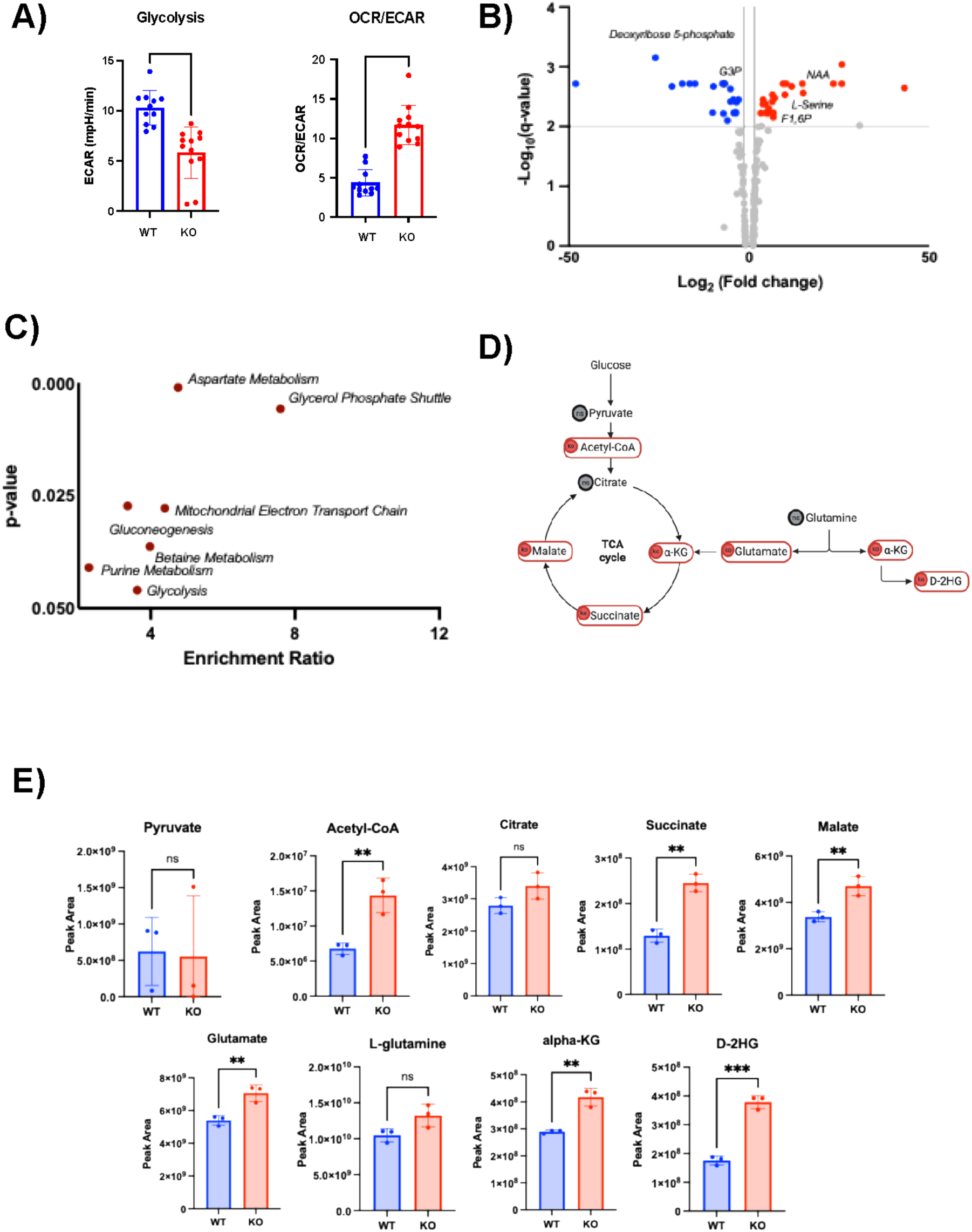
Lamin A/C expression modulates different energetic phenotypes in EBV+ B cells. **A)** Extracellular acidification rate (ECAR) and oxygen consumption rate (OCR) values measured by Seahorse analysis in WT and lamin A/C KO cells. The values obtained were used to calculate the OCR/ECAR ratio. Eleven independent measurements were performed. Data are expressed as mean ± SD (n = 11; p < 0.0001). **B)** Volcano plot of the metabolites differentially expressed in KO cells compared with WT cells. Significance defined as log_2_(|fold change|) > 1 and-log_10_(p-value) > 2.Upregulated metabolites are depicted in red, and downregulated genes are depicted in blue. **C)** Metabolite set enrichment analysis (MSEA) of the metabolic pathways differentially enriched between WT and lamin A/C K/O cells performed with MetaboAnalyst. Data are represented as a function of the enrichment ratio and p-value. **D)** Schematic representation of the changes in the TCA cycle observed in lamin A/C KO cells by metabolomics analysis. Metabolites with steady-state levels higher in KO cells compared with WT cells are circled in red; grey circles indicate no significant changes in the steady-state levels between conditions. **E)** Steady-state levels of the metabolites represented in D). Values represent the protein-normalized LC-MS peak areas for 3 biological samples for WT (blue) and KO (red) cell lines. All graphed data represent the mean ± SD of 3 biological replicates. *, p ≤ 0.05; **, p ≤ 0.01; ***, p ≤ 0.001; and ****, p ≤ 0.0001.

### Depletion of lamin A/C is linked to an increase in one-carbon metabolism and a reconfiguration of the amino acid pool in B cells

We hypothesized that lamin A/C depletion may redirect the glucose carbon flux into distinct metabolic pathways. To test this, we performed ^13^C-labeled glucose isotope tracing analysis to trace glucose-derived carbon in lamin A/C KO and WT B cells. Although both cell lines showed complete ^13^C-label incorporation for 3-phosphoglycerate (3PG, **Fig. 7A, B**), a glycolytic intermediate fundamental for the serine-glycine synthesis, we observed ^13^C incorporation for serine and glycine only in KO cells, indicating that in the absence of lamin A/C, cells increase the utilization of glucose for the SGOC pathway (**Fig. 7A, B**), which is particularly important during B-cell activation and germinal center formation and has been implicated in B-cell lymphomagenesis (117,121,122). We observed a significant increase in ^13^C incorporation for various other metabolites critical for one-carbon metabolism, including S-adenosylmethionine, in KO cells (**Fig. 7C, D**). Another interesting example of differential glucose flux was observed for the metabolism of N-acetyl-aspartate (NAA) and N-acetyl-aspartyl-glutamate (NAAG), 2 metabolites involved in alanine, aspartate, and glutamate metabolism (**Fig. S7B**). Carbon tracing revealed a higher proportion of labeled isotopologues in NAA from KO compared to the WT cells (39% vs 1% labeled), indicating increased incorporation of glucose-derived carbons in the synthesis of this metabolite, which is known as an acetate reservoir with roles in mitochondrial energy balance (**Fig. S7A**). In contrast, NAAG, a neuromodulator involved in glutamate regulation, exhibited more ^13^C labeling in WT cells (50%) than in KO cells (33%), suggesting that glutamate metabolism might be partially disrupted in the absence of lamin A/C expression (**Fig. S7A**) (123–126). This metabolic switch supports a regulatory role for lamin A/C in maintaining glutamate and acetyl-CoA homeostasis. In fact, the labeled fraction of acetyl-CoA was higher in WT cells (50%) than in KO cells (29%) (**Fig. S7A)**. At first glance, this may appear inconsistent with earlier data; however, as a major regulator of energy metabolism, molecular biosynthesis, and epigenetic modification, acetyl-CoA occupies a central position in cellular metabolism (127,128), therefore its labeling reflects input from many glucose-dependent pathways—not only the NAA/NAAG synthetic route. The NAA–NAAG pathway is only one of several contributors to its labeled pool, which helps explain why the labeling pattern does not directly mirror changes observed specifically in NAA/NAAG metabolism. Together, these results highlight a critical role for lamin A/C in regulating cellular metabolism in B cells. Its depletion promotes metabolic reprogramming that favors glycolysis-and TCA-derived amino acid biosynthesis, potentially supporting a proliferative and survival phenotype.

**Figure 7:**
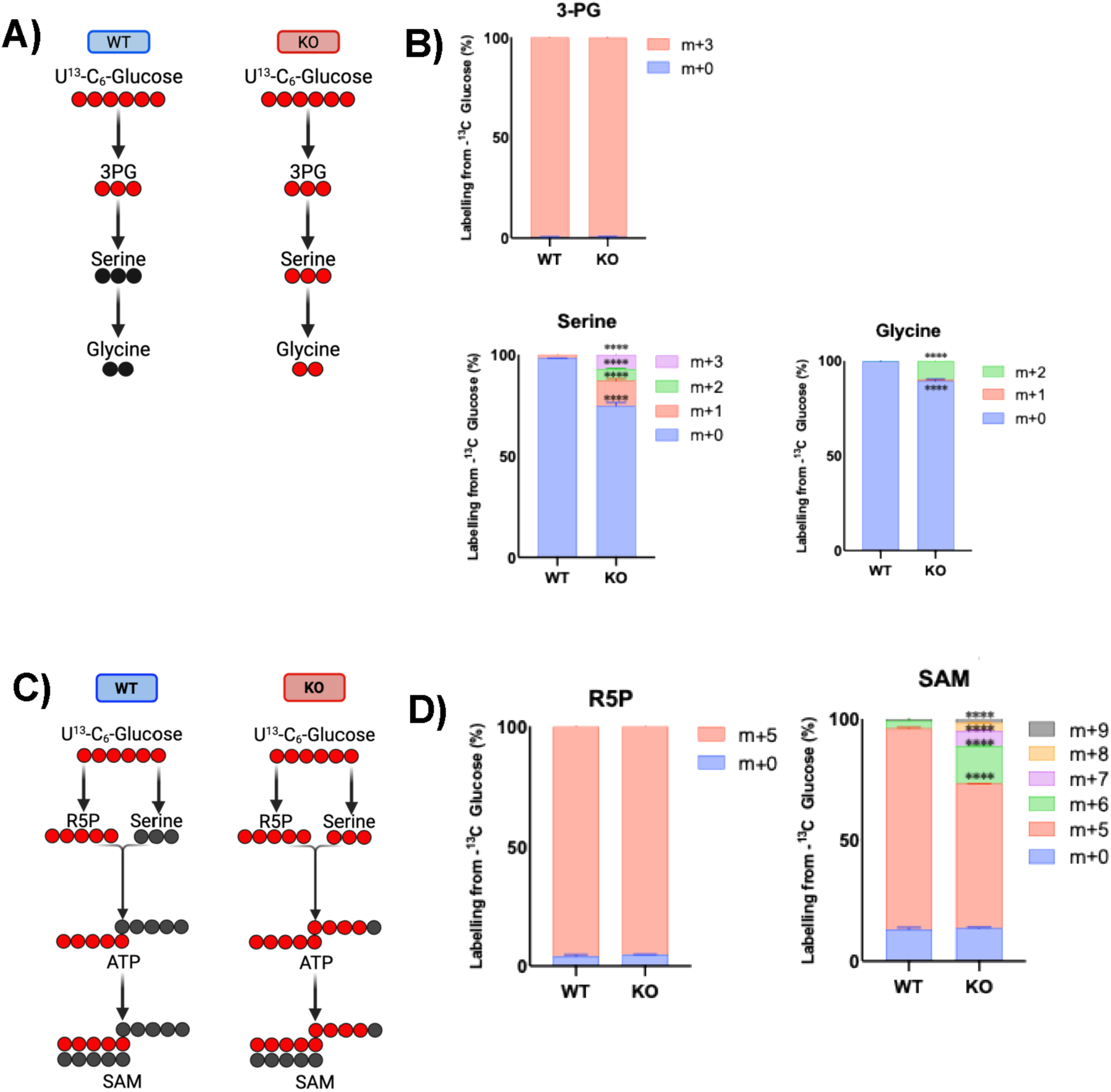
Lamin A/C depletion is linked to a redirection of glucose flux. **A)** Schematic illustration of carbon flux from glucose in the initial steps of the serine synthesis pathway in both WT and lamin A/C KO cells. ^13^C and ^12^C are represented with red and black circles, respectively. **B)** Isotope tracing data showing the ^13^C isotopologue distribution of polar metabolites extracted from WT and lamin A/C KO cells, corresponding to the metabolites in A). **C)** Schematic illustration of traced ^13^C-glucose in specific steps of the methionine cycle in both WT and lamin A/C KO cells. ^13^C and ^12^C are represented with red and black circles, respectively. **D)** Isotope tracing data showing the isotopologue distribution of polar metabolites extracted from WT and lamin A/C KO cells, corresponding to some of the metabolites in C): ribose-5-phosphate (R5P) and S-adenosylmethionine (SAM). All data in B) and D) are expressed as mean ± SD of 3 biological replicates. *, p ≤ 0.05; **, p ≤ 0.01; ***, p ≤ 0.001; and ****, p ≤ 0.0001.

### Lamin A/C depletion sensitizes B cells to PI3K inhibition

Our transcriptomic data, combined with the metabolomic studies, showed that lamin A/C is critical for regulating cellular pathways necessary for B-cell biology. We therefore hypothesized that depletion of lamin A/C in EBV+ B cells may result in vulnerabilities that could be targeted for therapeutic purposes. To test this, we screened WT and lamin A/C KO cells for their response to cancer drugs using the SelleckChem anti-cancer library, which contained 3500 compounds. We assessed the viability of WT and lamin A/C KO cells after 72 hours of treatment with each compound. Several compounds showed higher toxicity in KO cells than in WT cells (**Fig. S8A**). These included PI3K inhibitors (**Fig. 8A**), consistent with activation of the PI3K-AKT signaling pathway observed in KO cells through transcriptomic analysis (**Fig. S2** and **Fig. 5**) and protein expression analysis (**Fig. S8B**). The PI3K-AKT signaling pathway is frequently activated in cancer (129–132), affecting several critical cellular processes, including key metabolic pathways. We therefore decided to further investigate the effect of the well-known inhibitor acalisib (133), which we identified in our screening as one of the most toxic compounds in KO cells (**Fig. 8A**). Acalisib showed EC_50_ values of 0.003 µM in lamin A/C KO cells and 0.1 µM in WT cells, indicating that this drug is approximately 33 times more potent in lamin A/C-depleted cells (**Fig. 8B**). To confirm these findings, we assessed cell viability in WT and KO cells before and after treatment with 1 µM acalisib for 72 hours. Following treatment, only 25% of KO cells remained viable, compared with 75% of WT cells (**Fig. 8C**). Our data demonstrated that pharmacological inhibition of the H3K9-specific methyltransferase G9a in WT cells phenocopied the transcriptional effects observed upon lamin A/C depletion (**Fig. 5**). Notably, the PI3K signaling pathway was among the pathways similarly affected by both lamin A/C loss and G9a inhibition (**Fig. 5G**). Based on this overlap, we tested whether G9a inhibition sensitizes WT cells to acalisib. Consistent with this hypothesis, the combined treatment of WT cells with acalisib (0.1 µM) and a G9a inhibitor (1 µM) for 72 hours resulted in a significant reduction in cell viability compared with untreated controls and cells treated with the G9a inhibitor alone (**Fig. 8D**). G9a inhibition alone did not impact cell viability but enhanced sensitivity to PI3K inhibition. Together, these data suggest that lamin A/C regulates PI3K pathway activity, and its depletion renders B cells more sensitive to PI3K inhibition. In cells expressing lamin A/C, inhibition of H3K9me2 deposition can mimic the epigenetic landscape of lamin A/C-deficient cells, thereby sensitizing them to PI3K inhibition.

**Figure 8:**
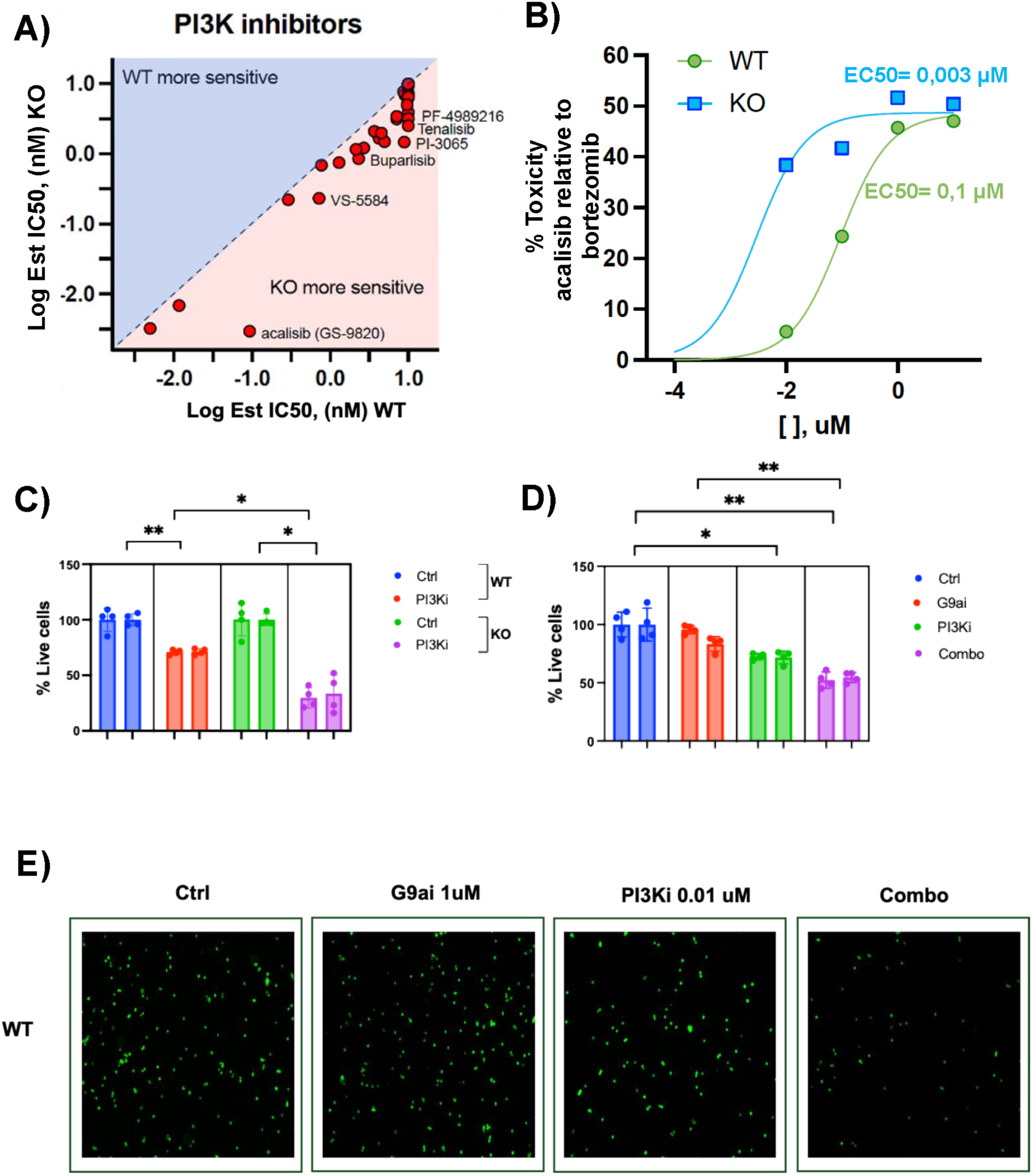
Lamin A/C depletion sensitizes B cells to PI3K inhibition. **A)** Drug screening assay on WT and lamin A/C KO cells treated with a library of PI3K inhibitors. Each compound was tested at 10, 1, 0.1, and 0.01 μM. Cell viability was assessed by measuring luminescence and converting the luminescence values to % toxicity. The graph shows the correlation between the log IC_50_ in WT and KO cells for each compound. **B)** Acalisib dose-response curve in WT and lamin A/C KO cells. Toxicity was calculated using a luminescence assay, where the luminescence values were converted to % toxicity. Treatment with bortezomib (1 μM) was included as a positive control (100% toxicity); 0% toxicity corresponded to the RLU values of negative controls (0.1% DMSO). IC_50_ values for acalisib in WT and lamin A/C KO cells are indicated. **C)** Cell viability (% live cells) assessed by counting the number of live cells interacting with the green dye calcein AM in WT and lamin A/C KO cells after treating for 72 hours with the PI3K inhibitor (PI3Ki) acalisib (0.1 µM) or control (Ctrl, 0.1% DMSO). Data come from 2 experiments with 4 biological replicates each (*, p ≤ 0.05; **, p ≤ 0.01). Sample acquisition and analysis were conducted using a fluorescence microscope. Fluorescence was quantified using the ImageJ software. **D)** Cell viability (% live cells) assessed by counting the number of live WT cells interacting with calcein AM in samples treated with control (Ctrl, 0.1% DMSO), a G9a inhibitor (G9ai, 1 µM), acalisib (PI3Ki, 0.01 µM), or their combination. Data come from 2 experiments with 4 biological replicates each (*, p ≤ 0.05; **, p ≤ 0.01) Sample acquisition and analysis were conducted using a fluorescence microscope. Fluorescence was quantified using the ImageJ software. **E)** Representative fluorescence microscope images of WT cells treated for 72 hours with control (Ctrl, 0.1% DMSO), G9a inhibitor (1 µM), acalisib (0.01 µM), or their combination and stained using calcein AM.

## DISCUSSION

Our study reveals a novel and crucial role for lamin A/C in reorganizing the nuclear space and regulating host gene expression in EBV-infected B cells, with broad implications for understanding how EBV may manipulate host nuclear architecture to support its lifecycle and influence B-cell fate (134–136). The role of lamin A/C in gene regulation has been well studied due to its nuclear localization and ability to interact with key transcriptional regulators. Lamin A/C is associated with maintaining nuclear structure and organizing chromatin by forming lamina-associated domains (LADs) (103). In the context of viral infection, we previously showed that EBV infection results in increased expression of host lamin A/C and that the latter binds to and controls the viral genome expression (56). However, whether EBV reorganizes the nuclear peripheral space to regulate host gene expression remains unclear, and the mechanisms by which lamin A/C influences gene expression and causes transcriptional reprogramming in B cells are still unknown. Here, we present evidence that lamin A/C actively and dynamically participates in modulating transcriptional programs crucial for B-cell differentiation, metabolism, and proliferation through altering 3D genome organization, heterochromatin deposition, and by affecting CTCF nuclear localization.

We demonstrate that lamin A/C depletion in EBV-infected B cells leads to broad transcriptional reprogramming affecting key pathways involved in B-cell biology, such as PI3K signaling, the UPR pathway, and actin cytoskeleton regulation via the ARP2/3-WASP complex (83,84). These pathways are integral to B-cell activation and transformation, suggesting that lamin A/C expression may be essential to create a transcriptional environment favorable for B-cell proliferation. Our transcriptomic data showed that lamin A/C KO cells present a cellular phenotype similar to naïve B-cells, indicating a possible role of lamin A/C in B-cell activation and differentiation. Interestingly, we also observed that, in this model, ablation of lamin A/C expression resulted in the transcriptional inhibition of key metabolic pathways, including glycolysis and gluconeogenesis, which was supported by mass spectrometry metabolic data.

Lamin A/C KO cells showed enrichment in pathways related to energy production and biosynthesis of macromolecules such as nucleotides, lipids, and proteins. Notably, the TCA cycle, one-carbon metabolism, and the pentose phosphate pathway exhibited increased levels of various metabolites. These pathways are essential for B-cell biology and support the ability of the cells to respond flexibly to stimuli, transitioning from a resting to an activated, highly proliferative state, or differentiating into plasma or memory cells. EBV infection hijacks these pathways to regulate different latency programs. Previous studies (62,64,137) have demonstrated that EBV fundamentally rewires host cell metabolism to create a favorable environment for persistence and transformation of B cells into immortalized cells, notably inducing mitochondrial one-carbon metabolism and the expression of related enzymes. Our findings indicate that lamin A/C contributes to metabolic regulation, at least in part, by controlling the transcriptional landscape of key metabolic pathways, supporting a model in which lamin A/C helps sustain the metabolic state necessary for efficient viral-driven B-cell proliferation and differentiation. Our data suggest that EBV exploits lamin A/C–dependent chromatin and transcriptional programs to secure the metabolic conditions required for successful infection and transformation.

Antigen-driven and virus-induced B-cell activation are both associated with extensive remodeling of the genome 3D architecture (135,138). Our data indicate that lamin A/C expression plays a significant role in this process, as its depletion produced a marked shift in 3D genome organization, characterized by a loss of long-range chromatin loops and a compensatory increase in short-range contacts. This reorganization suggests that lamin A/C is required to maintain extended regulatory interactions that coordinate transcription across large genomic distances. The reduction of long-range loops likely disrupts enhancer–promoter communication and higher-order transcriptional networks, while the enrichment of short-range interactions reflects a collapse of large-scale chromatin architecture into more locally compacted domains. Lamin A/C-mediated topological changes are functionally relevant, as the loss or gain of specific chromatin loops correlated with transcriptional changes. Together, these observations indicate that lamin A/C is essential for preserving higher-order genome topology, and its absence leads to fragmentation of chromatin architecture with consequences for gene regulation and cellular function (139–141).

Our data suggest that lamin A/C–dependent control of higher-order genome topology may be mediated, at least in part, by its interaction with CTCF. Although lamin A/C depletion did not alter CTCF occupancy across the genome, CTCF-bound genes were preferentially downregulated in lamin A/C KO cells and showed increased accumulation of the peripheral heterochromatin mark H3K9me2 (51,54,142). Prior work has established that H3K9me2-enriched regions preferentially localize to the nuclear periphery (111)—a repressive compartment —consistent with the notion that lamin A/C helps maintain proper nuclear positioning of CTCF-regulated loci. In line with this model, our PLA assays revealed that in the absence of lamin A/C, CTCF displayed enhanced interaction with lamin B1, therefore a more peripheral distribution, relative to WT cells. Such relocalization likely limited the ability of CTCF to support long-range chromatin looping, thereby contributing to the collapse of genome architecture and the transcriptional dysregulation observed in lamin A/C KO cells. Together, these observations support a model in which CTCF-bound loci dynamically oscillate between the nuclear interior and periphery depending on lamin A/C levels. Upon antigen-driven or EBV-induced B-cell activation, lamin A/C may stabilize a central nuclear positioning of CTCF-regulated chromatin domains, allowing these regions to engage in extensive long-range interactions and support active transcriptional programs. Conversely, in the absence of lamin A/C, CTCF-bound loci may drift toward the nuclear periphery, where increased association with lamin B1 and H3K9me2 deposition promote a more repressive environment. Although further work will be needed to resolve the dynamics of this proposed “positional oscillation,” this model offers a framework linking lamin A/C expression, nuclear compartmentalization, and CTCF-mediated transcriptional regulation.

This proposed model is further supported by our finding that blocking H3K9me2 deposition phenocopies the transcriptional profile of lamin A/C depletion. In WT B cells, inhibition of the histone methyltransferase G9a, which catalyzes H3K9me2 deposition at LADs, altered the global transcriptome similarly to lamin A/C depletion (143–145). IPA analysis of the common targets between lamin A/C KO and histone methyltransferase G9a inhibition revealed consistent changes in signaling and metabolic pathways, including inhibition of glycolysis/gluconeogenesis and activation of PI3K/AKT and ATM signaling. These results support a collaborative and dynamic role for lamin A/C and H3K9me2 in maintaining transcriptional programs relevant to B-cell function and metabolism and align with other studies showing that loss of lamin A/C causes the heterochromatin to be untethered to the nuclear periphery, disrupting its supporting role (146,147).

Given the importance of lamin A/C for B-cell activation and EBV-driven proliferation, the phenocopy effect of G9a blockade on global transcription has potential translational implications, particularly regarding the link with the PI3K pathway, which is fundamental for B-cell differentiation, survival, and immunological function (83,84). This pathway is also targeted by EBV to support viral persistence and survival in the host environment. PI3K inhibitors have been tested for the treatment of B-cell malignancies with good results, although in the case of EBV-induced cancer, the use of PI3K inhibitors can induce drug resistance (148,149). Consistent with the observed upregulation of PI3K-AKT pathway components, lamin A/C KO cells showed heightened sensitivity to PI3K inhibitors, including acalisib. Interestingly, in our model, G9a inhibition sensitized WT cells to acalisib, further suggesting that disruption of H3K9 methylation can phenocopy lamin A/C loss and expose similar therapeutic vulnerabilities. In future work, we plan to determine the synergistic effect of G9a inhibitors and acalisib on tumor growth in a preclinical model of EBV-driven malignancies.

One limitation of the present study is the reliance on a single EBV-positive B-cell line model, which may not fully recapitulate in vivo conditions. Future research should expand these analyses to additional B-cell models and, ideally, to primary human samples from individuals with EBV-associated diseases or laminopathies. Moreover, our conclusions are based on observations conducted on a lamin A/C knockout cell line. While this model is highly useful for dissecting the role of lamin A/C in host–virus interactions, it partially lacks insights into how lamin A/C post-translational modifications (PTMs)—such as phosphorylation—contribute to this process. This represents an important gap, as lamin A/C PTMs are critical regulators of lamin function (150,151), and their dysregulation is linked to structural alterations and susceptibility to various diseases and infections (152–154). Although our previous work demonstrated that EBV infection is sufficient to induce lamin A/C expression, and this present work establishes the importance of lamin A/C in supporting transcriptional changes necessary for EBV-driven B-cell proliferation, the specific EBV protein responsible for inducing lamin A/C activation remains unidentified. Further investigation is needed to elucidate the viral factors and molecular mechanisms that regulate lamin A/C during EBV infection.

Together, our findings establish lamin A/C as a key regulator of nuclear architecture and gene expression, operating through both structural mechanisms—such as controlling higher-order chromatin topology—and biochemical mechanisms, including the modulation of epigenetic states. Numerous other studies have demonstrated a strong relationship between lamin A/C and DNA virus infection. Lamin–virus interactions can occur through multiple routes: lamin A/C may serve as a scaffold that supports viral genome replication or transcription, or it may influence viral nuclear egress (57,155,156). Also, many DNA viruses encode proteins capable of disrupting lamin organization or activating signaling pathways that alter lamin structure, stability, or function, thereby facilitating viral entry or replication (157). Collectively, these studies define lamin A/C as an important antiviral barrier, particularly during the lytic phase of infection. In the context of EBV infection, lamin A/C appears to play a dual role—contributing to host antiviral defense during lytic replication, while being hijacked by EBV during latency to mediate remodeling of host chromatin, promote viral persistence, and reprogram B-cell biology by reshaping transcriptional and metabolic pathways.

## DATA AVAILABILITY

The RNA-seq, the ChIP-seq and Hi-C datasets generate during this study are available in the Gene Expression Omnibus (GEO) repository, under accession number GSE314558.

## ACKNOWLEDGEMENTS

We are grateful to The Wistar Institute’s Proteomics and Metabolomics Shared Resource for providing technical support.

## AUTHOR CONTRIBUTIONS

Conceptualization: L.B.C, I. T.; Data curation: L.B.C., Sa.S., S.S., A.K., A.R.G., J.C.; Formal analysis: L.B.C., D.M., S.S., A.K.; Funding acquisition: I.T.; Investigation: L.B.C.; Project administration: I.T.; Supervision: I.T.; Validation: L.B.C.; Visualization: L.B.C., I.T.; Writing original draft: L.B.C, I.T.; Writing – review & editing: L.B.C, I.T., A.R.G.

