## Supplemental figures for "Lamin A/C maintains genome topology and regulates transcriptional programs essential for virus-driven B cell activation"

**Supplemental Figure 1**

### Figure S1

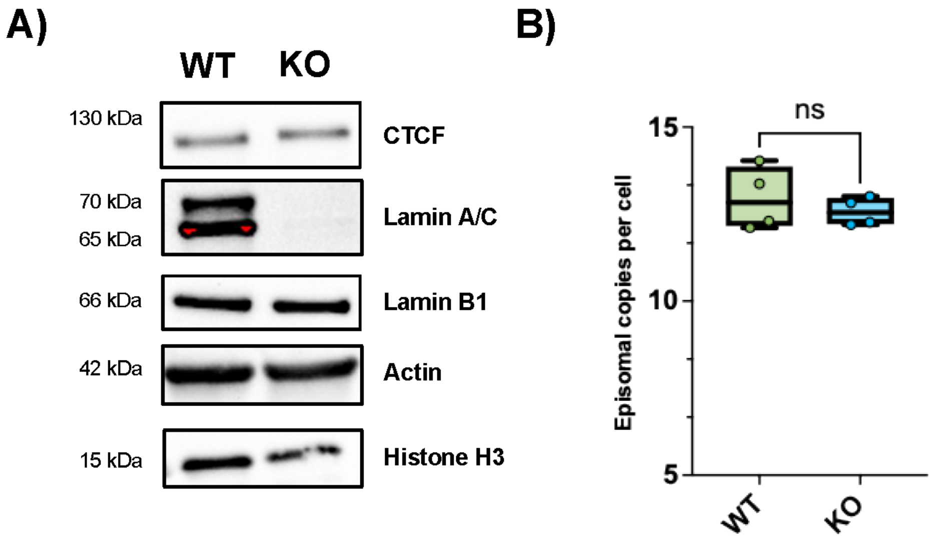


**Figure S1: A)** Western blot showing expression of CTCF, lamin A/C, lamin B1, and housekeeping controls (actin and histone H3) in WT and lamin A/C KO cells. **B)** Digital droplets PCR (ddPCR) measuring EBV episomal copy number per cell in WT and lamin A/C KO cell lines. Four different samples were analyzed, no significant changes were detected.

**Supplemental Figure S2**

### Figure S2

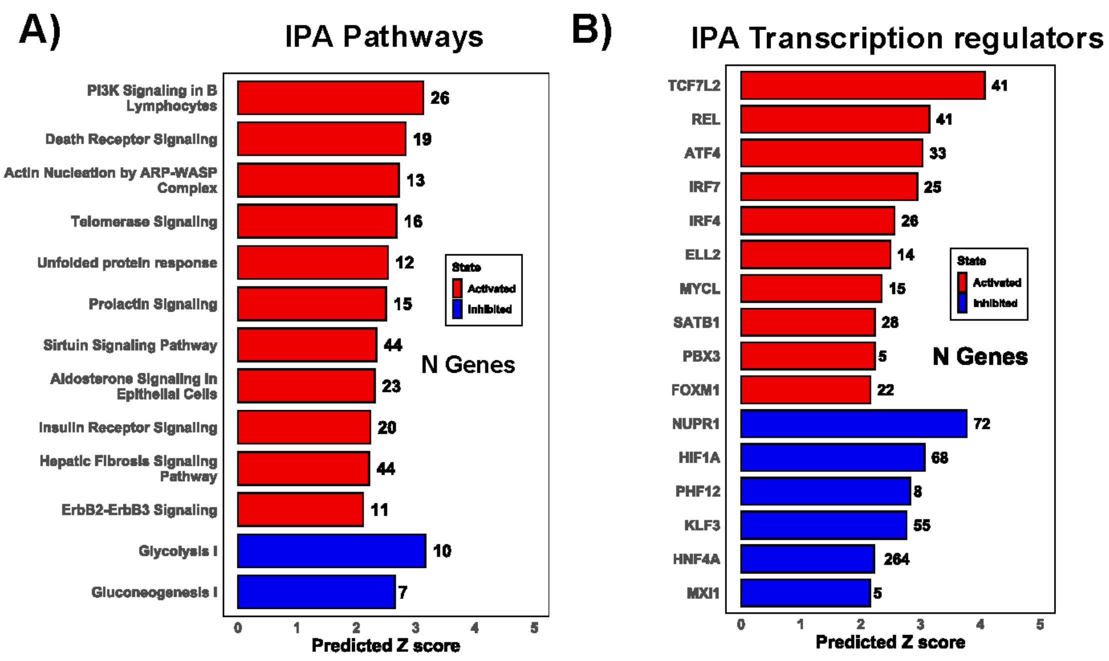


**Figure S2: A)** Ingenuity pathway analysis (IPA) of the top 13 pathways significantly affected by lamin A/C KO in LCL cells, as determined by RNA-Seq. **B)** IPA analysis showing the top 10 transcriptional regulators activated and top 6 inhibited by lamin A/C KO. Predicted z scores and the number of genes relative to each of the pathways are indicated in the graphs.

#### Supplemental Figure 3

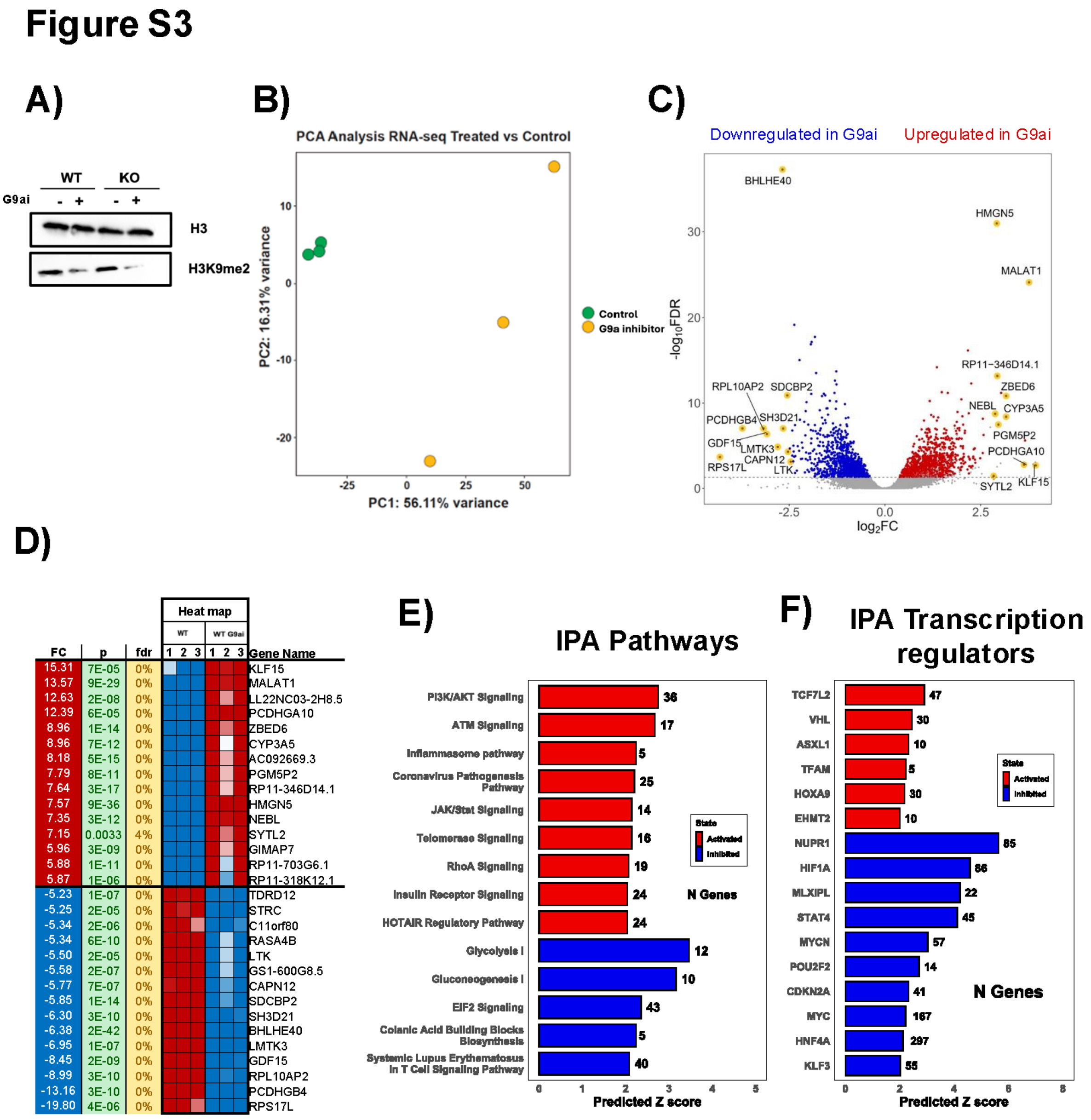


**Figure S3: A)** Western blot showing H3K9me2 levels in WT and lamin A/C KO cells treated with the G9a inhibitor UNC0631 (1 µM) for 72h. H3 is shown as housekeeping control. **B)** Principal component analysis (PCA) scatter plot of RNA-Seq data from WT cells treated with the G9a inhibitor and control. The samples are shown as a function of principal component 1, or PC1 (x-axis), and principal component 2, or PC2 (y-axis). The percentage of variance accounted for by PC1 and PC2 is indicated**.** **C)** Volcano plot of the 2,346 differentially expressed genes (false discovery rate (FDR) < 5%, $|\log_2$ fold change (FC)$| > 1.5$) in WT cells treated with G9a inhibitor (G9ai) compared with WT control. Upregulated genes are depicted in red, and downregulated genes in blue. Genes with the highest change in expression are highlighted in yellow. **D)** Heat map of RNA-Seq data showing the top 30 genes whose expression was significantly altered (FRD < 5%) by G9ai treatment compared to WT control cells. p-values are indicated. **E)** Ingenuity Pathway Analysis (IPA) of the top canonical pathways significantly affected by G9ai treatment in WT cells, as determined by RNA-Seq. **F)** IPA showing the top upstream transcriptional regulators affected by G9a inhibition in WT cells.

Predicted z scores and the number of genes relative to each of the pathways are indicated in the graphs.

#### Supplemental Figure 4

### Figure S4

##### Volcano plot of 1857 genes that overlap between G9ai/Control in WT and KO/WT in control

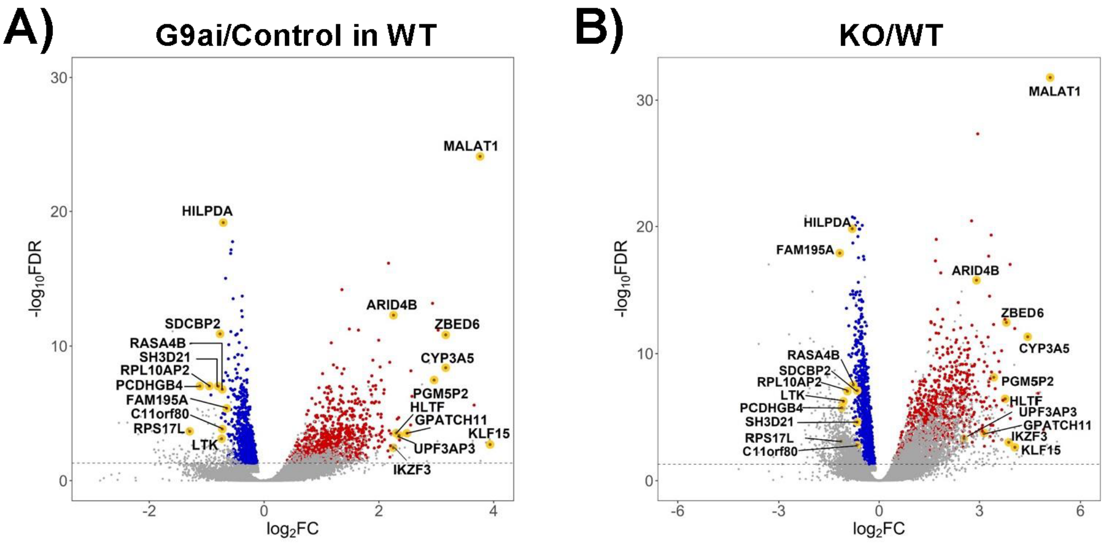


C)

| Fold: WT G9Ai vs WT ct | Fold: KO ct vs WT ct | p: WT G9Ai vs WT ct | p: KO ct vs WT ct | FDR: WT G9Ai vs WT ct | FDR: KO ct vs WT ct | Heatmap WT ct 1 | Heatmap WT ct 2 | Heatmap WT ct 3 | Heatmap WT G9Ai 1 | Heatmap WT G9Ai 2 | Heatmap WT G9Ai 3 | Heatmap KO 2 | Heatmap KO 3 | Gene |
|---|---|---|---|---|---|---|---|---|---|---|---|---|---|---|
| 15.31 | 16.66 | 7E-05 | 0.0003 | 0% | 0% | | | | | | | | | KLF15 |
| 13.57 | 34.14 | 9E-29 | 7E-37 | 0% | 0% | | | | | | | | | MALAT1 |
| 12.63 | 26.60 | 2E-08 | 3E-09 | 0% | 0% | | | | | | | | | LL22NC03-2H8.5 |
| 8.96 | 13.98 | 1E-14 | 1E-15 | 0% | 0% | | | | | | | | | ZBED6 |
| 8.96 | 21.75 | 7E-12 | 2E-14 | 0% | 0% | | | | | | | | | CYP3A5 |
| 8.18 | 13.56 | 5E-15 | 5E-16 | 0% | 0% | | | | | | | | | AC092669.3 |
| 7.79 | 10.76 | 8E-11 | 9E-11 | 0% | 0% | | | | | | | | | PGM5P2 |
| 7.64 | 15.09 | 3E-17 | 9E-21 | 0% | 0% | | | | | | | | | RP11-346D14.1 |
| 5.88 | 10.87 | 1E-11 | 6E-13 | 0% | 0% | | | | | | | | | RP11-703G6.1 |
| 5.87 | 15.54 | 1E-06 | 4E-08 | 0% | 0% | | | | | | | | | RP11-318K12.1 |
| 5.61 | 8.81 | 6E-06 | 1E-05 | 0% | 0% | | | | | | | | | GPATCH11 |
| 5.10 | 5.79 | 1E-05 | 4E-05 | 0% | 0% | | | | | | | | | UPF3AP3 |
| 5.08 | 9.21 | 2E-07 | 7E-09 | 0% | 0% | | | | | | | | | AC008394.1 |
| 4.93 | 13.65 | 6E-06 | 9E-09 | 0% | 0% | | | | | | | | | HLTF |
| 4.77 | 7.52 | 3E-16 | 2E-19 | 0% | 0% | | | | | | | | | ARID4B |
| 4.76 | 14.67 | 0.0002 | 0.0001 | 0% | 0% | | | | | | | | | IKZF3 |
| 4.99 | 4.75 | 3E-07 | 4E-06 | 0% | 0% | | | | | | | | | CHN2 |
| 4.62 | 11.35 | 7E-06 | 4E-08 | 0% | 0% | | | | | | | | | GOLGA4 |
| 4.60 | 15.16 | 1E-04 | 6E-07 | 0% | 0% | | | | | | | | | RB1CC1 |
| 4.57 | 29.70 | 0.0013 | 7E-06 | 2% | 0% | | | | | | | | | ALG11 |
| 4.56 | 7.51 | 2E-12 | 2E-14 | 0% | 0% | | | | | | | | | ANKRD12 |
| 4.53 | 13.01 | 4E-06 | 1E-08 | 0% | 0% | | | | | | | | | DPY19L2P4 |
| 4.49 | 6.79 | 2E-20 | 9E-25 | 0% | 0% | | | | | | | | | HMGB2 |
| 4.46 | 12.69 | 0.0003 | 6E-05 | 1% | 0% | | | | | | | | | TTC32 |
| 4.28 | 6.51 | 3E-05 | 1E-05 | 0% | 0% | | | | | | | | | SPATA13 |
| 4.26 | 5.15 | 7E-10 | 8E-10 | 0% | 0% | | | | | | | | | IFT57 |
| 4.24 | 7.01 | 1E-09 | 5E-11 | 0% | 0% | | | | | | | | | SWAP70 |
| 4.17 | 12.18 | 4E-06 | 3E-11 | 0% | 0% | | | | | | | | | KRCC1 |
| 4.16 | 11.91 | 1E-06 | 3E-12 | 0% | 0% | | | | | | | | | MORC4 |
| 4.12 | 9.28 | 1E-05 | 7E-08 | 0% | 0% | | | | | | | | | RP11-386D6.1 |

D)

| Fold: WT G9Ai vs WT ct | Fold: KO ct vs WT ct | p: WT G9Ai vs WT ct | p: KO ct vs WT ct | FDR: WT G9Ai vs WT ct | FDR: KO ct vs WT ct | Heatmap WT ct 1 | Heatmap WT ct 2 | Heatmap WT ct 3 | Heatmap WT G9Ai 1 | Heatmap WT G9Ai 2 | Heatmap WT G9Ai 3 | Heatmap KO 2 | Heatmap KO 3 | Gene |
|---|---|---|---|---|---|---|---|---|---|---|---|---|---|---|
| -3.55 | -4.42 | 4E-22 | 1E-24 | 0% | 0% | | | | | | | | | SLX1A |
| -3.57 | -4.36 | 0.0038 | 0.001 | 4% | 1% | | | | | | | | | KRT223P |
| -3.59 | -8.09 | 3E-07 | 2E-15 | 0% | 0% | | | | | | | | | DLG4 |
| -4.25 | -3.67 | 0.0009 | 0.0085 | 1% | 4% | | | | | | | | | SERHL2 |
| -3.68 | -3.86 | 5E-07 | 7E-06 | 0% | 0% | | | | | | | | | GPR146 |
| -4.65 | -3.69 | 2E-11 | 3E-07 | 0% | 0% | | | | | | | | | RP11-475J5.8 |
| -8.45 | -3.73 | 2E-09 | 0.0004 | 0% | 0% | | | | | | | | | GDF15 |
| -3.76 | -4.25 | 3E-05 | 0.0002 | 0% | 0% | | | | | | | | | RPL23AP7 |
| -3.80 | -5.48 | 4E-08 | 2E-11 | 0% | 0% | | | | | | | | | ALDOC |
| -4.84 | -3.87 | 1E-07 | 3E-05 | 0% | 0% | | | | | | | | | UPK2 |
| -3.94 | -5.03 | 0.0007 | 6E-05 | 1% | 0% | | | | | | | | | FXYD2 |
| -4.00 | -4.98 | 8E-11 | 2E-11 | 0% | 0% | | | | | | | | | NUDT8 |
| -4.04 | -5.37 | 1E-05 | 3E-06 | 0% | 0% | | | | | | | | | RP11-268J15.5 |
| -4.69 | -4.10 | 3E-19 | 7E-14 | 0% | 0% | | | | | | | | | MT2A |
| -4.13 | -5.33 | 1E-13 | 3E-15 | 0% | 0% | | | | | | | | | LIPE |
| -6.38 | -4.14 | 2E-42 | 2E-21 | 0% | 0% | | | | | | | | | BHLHE40 |
| -4.16 | -7.43 | 9E-07 | 7E-11 | 0% | 0% | | | | | | | | | LIME1 |
| -5.58 | -4.24 | 2E-07 | 7E-05 | 0% | 0% | | | | | | | | | GS1-600G8.5 |
| -4.30 | -4.72 | 3E-11 | 2E-10 | 0% | 0% | | | | | | | | | LTBP3 |
| -4.49 | -15.04 | 3E-08 | 8E-22 | 0% | 0% | | | | | | | | | FAM195A |
| -5.85 | -4.51 | 1E-14 | 1E-09 | 0% | 0% | | | | | | | | | SDCBP2 |
| -6.30 | -4.53 | 3E-10 | 1E-06 | 0% | 0% | | | | | | | | | SH3D21 |
| -5.34 | -4.58 | 2E-06 | 0.0002 | 0% | 0% | | | | | | | | | C11orf80 |
| -5.09 | -4.78 | 1E-10 | 1E-08 | 0% | 0% | | | | | | | | | RP11-611D20.2 |
| -5.15 | -6.36 | 1E-23 | 6E-24 | 0% | 0% | | | | | | | | | HILPDA |
| -5.34 | -5.92 | 6E-10 | 4E-10 | 0% | 0% | | | | | | | | | RASA4B |
| -5.50 | -11.71 | 2E-05 | 1E-08 | 0% | 0% | | | | | | | | | LTK |
| -8.99 | -8.89 | 3E-10 | 2E-09 | 0% | 0% | | | | | | | | | RPL10AP2 |
| -13.16 | -13.16 | 3E-10 | 6E-08 | 0% | 0% | | | | | | | | | PCDHGB4 |
| -19.80 | -13.86 | 4E-06 | 8E-05 | 0% | 0% | | | | | | | | | RPS17L |

Fold: ≤ -2, -1.5, -1.2, -1.1, ±1, 1.1, 1.2, 1.5, ≥ 2

**Figure S4**: **A)** and **B)** Volcano plots of the 1,857 genes that are regulated by G9a inhibitor treatment in WT cells (A) and by lamin A/C KO (B). Upregulated genes are depicted in red, and downregulated genes in blue. Genes with the highest change in expression are highlighted in yellow. **C)** and **D)** Heatmaps of RNA-Seq data showing the top targets whose expression was significantly altered (FRD < 5%) and that are concordantly upregulated (C) and downregulated (D) by lamin A/C KO and G9a inhibitor (G9ai) treatment. Fold changes, false discovery rate (FDR) values, and p-values are shown.

**Supplemental Figure 5**

### Figure S5

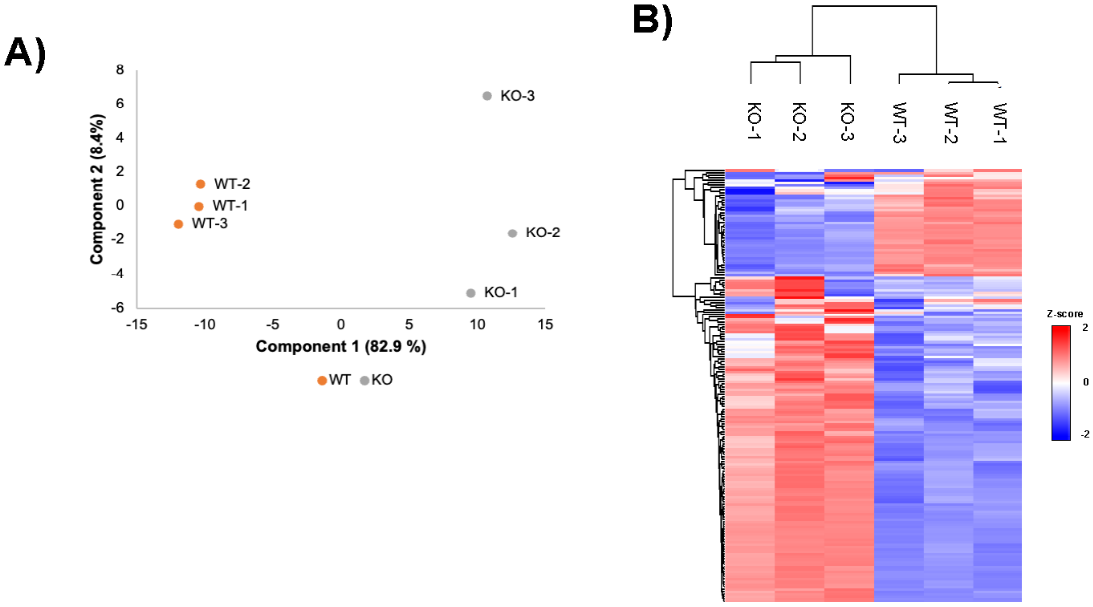


**Figure S5: A)** Principal component analysis (PCA) scatter plot of mass spectrometry metabolic data from WT and lamin A/C KO cells, showing that the 2 cell lines cluster differently based on their metabolic expression profile. The samples are shown as a function of principal component 1, or PC1 (x-axis), and principal component 2, or PC2 (y-axis). The percentage of variance accounted for by PC1 and PC2 is indicated. **B)** Heat map of mass spectrometry metabolic data showing differentially expressed metabolites in WT and lamin A/C KO cells. Color code represents different z-score values, with upregulated and downregulated metabolites shown in red and blue, respectively.

**Supplemental Figure 6**

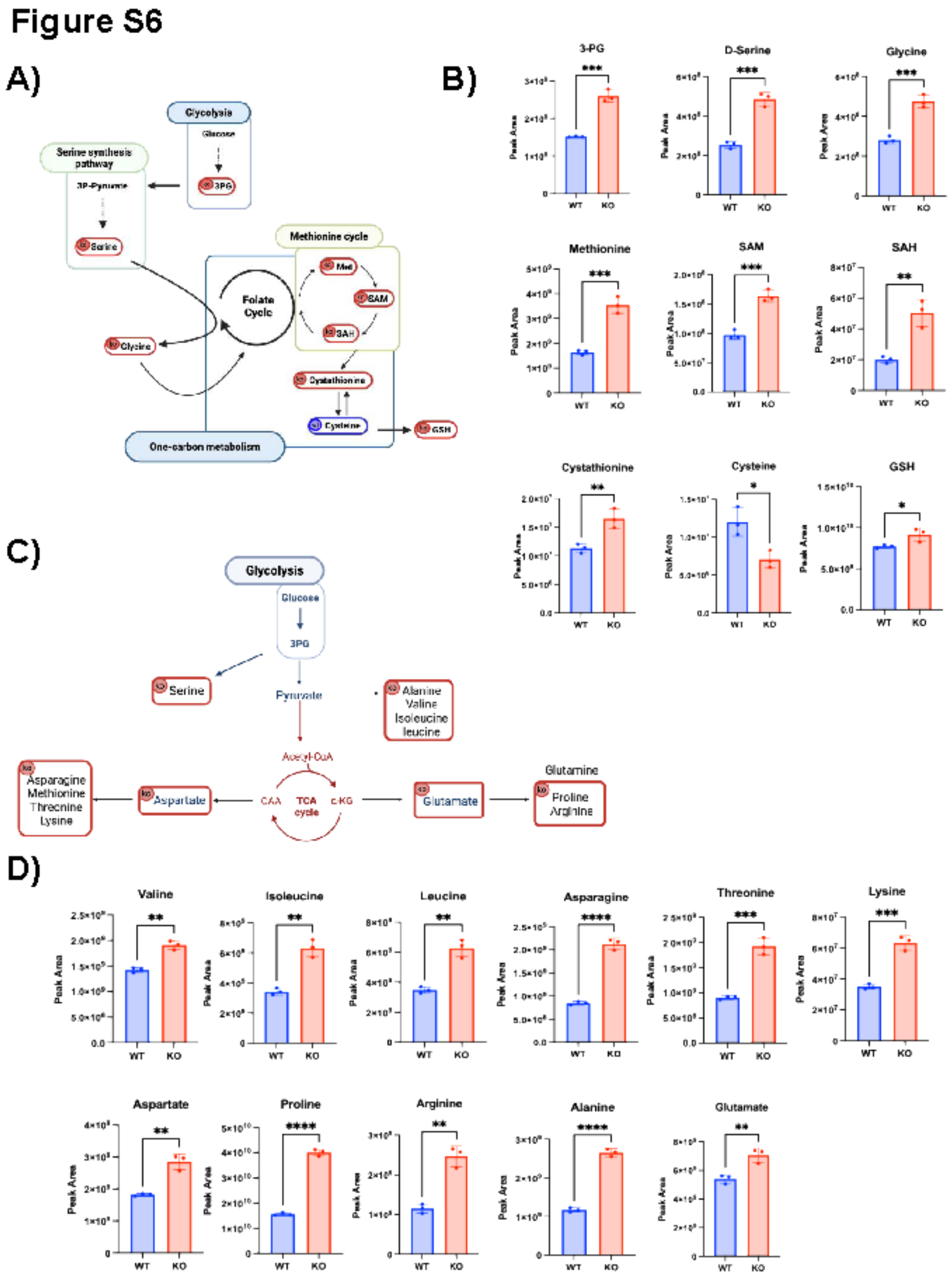


**Figure S6: A)** Schematic representation of the changes in the glycine–serine and one-carbon pathway (SGOC) observed in lamin A/C KO cells by metabolomics analysis. Metabolites with steady-state levels that were significantly affected are circled in red (higher in KO cells) and blue (higher in WT cells). **B)** Steady-state levels of the metabolites represented in A). Values are the protein-normalized LC-MS peak areas for WT (blue) and KO (red) cell lines. **C)**

Schematic representation of the changes in the biosynthesis of amino acids observed in lamin A/C KO cells by metabolomics analysis. Metabolites with significantly higher steady-state levels in KO compared to WT cells are circled in red. **D)** Steady-state levels of the metabolites represented in C). Values are the protein-normalized LC-MS peak areas for WT (blue) and KO (red) cell lines. All graphed data in B) and D) represent the mean ± SD of 3 biological replicates. *, $p \leq 0.05$; **, $p \leq 0.01$; ***, $p \leq 0.001$, and ****, $p \leq 0.0001$.

**Supplemental Figure 7**

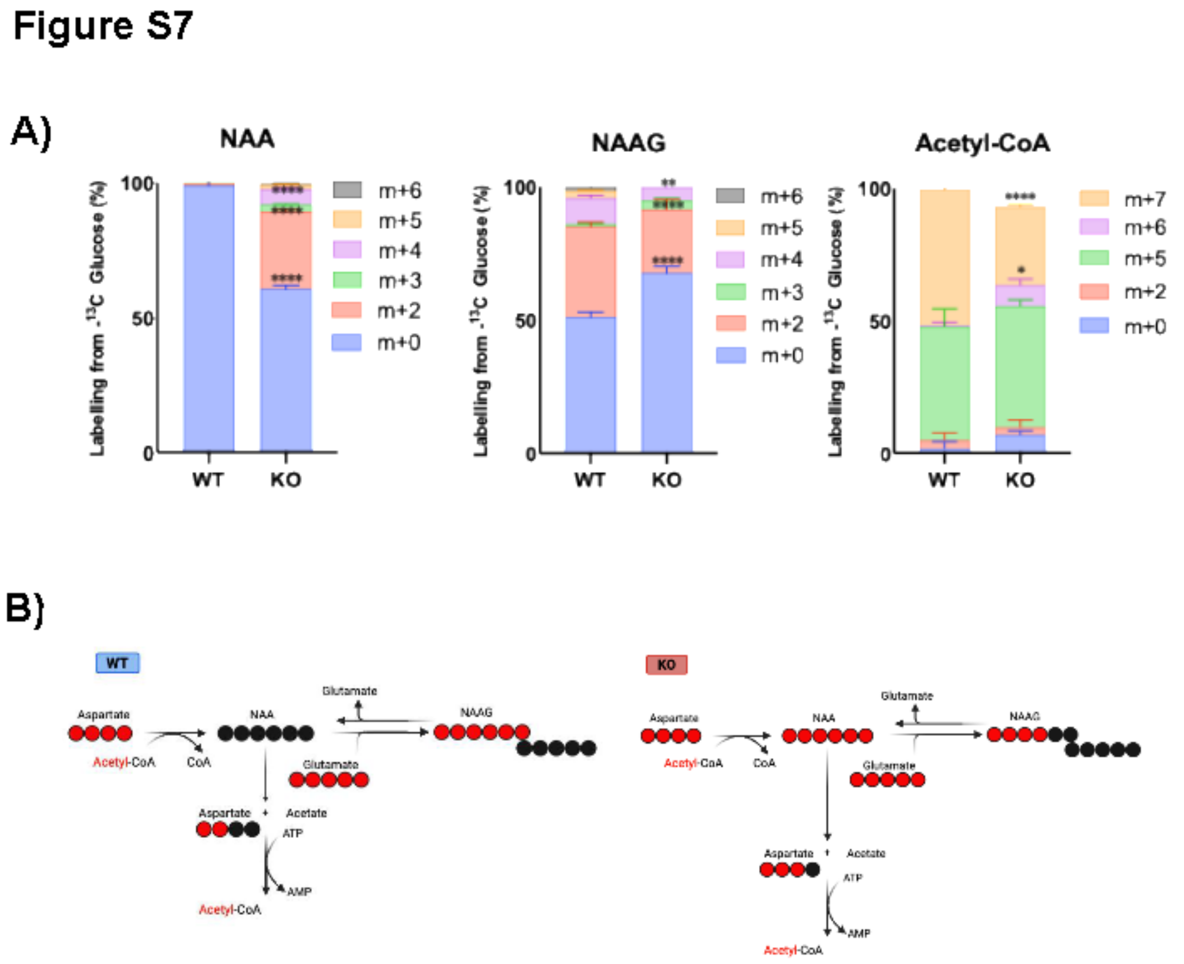


**Figure S7: A)** $^{13}C$ isotope tracing showing the isotopologue distributions of metabolites in the NAA-NAAG pathway in samples extracted from WT and lamin A/C KO cells. Data represent the mean ± SD of 3 biological replicates. *, p ≤ 0.05; **, p ≤ 0.01; ****, p ≤ 0.0001. NAA: N-acetylaspartate, NAAG: N-acetylaspartylglutamate. **B)** Schematic illustration of traced $^{13}C$-glucose in specific steps of the NAA-NAAG pathway in WT and lamin A/C KO cells. $^{13}C$ and $^{12}C$ are represented with red and black circles, respectively.

**Supplemental Figure 8**

### Figure S8

A)

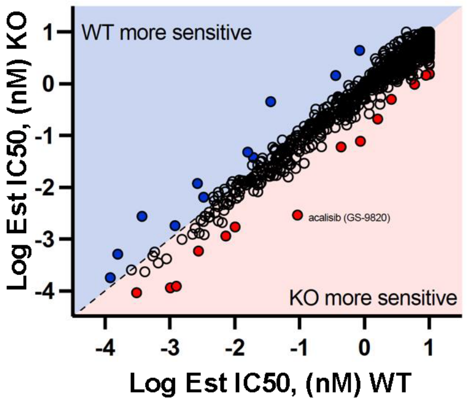


B)

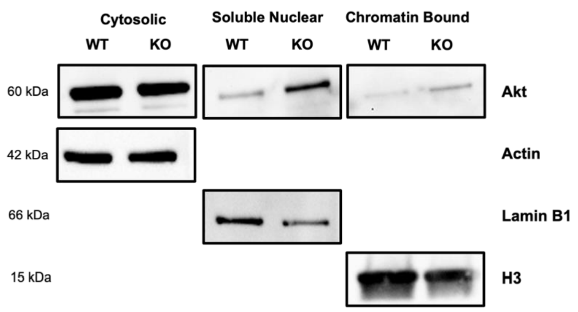


**Figure S8: A)** Drug screening assay on WT and lamin A/C KO cells treated with a library of anticancer drugs. Each compound was tested at 10, 1, 0.1, and 0.01 μM. Cell viability was assessed by measuring luminescence. The luminescence values were converted to % toxicity. The graph shows the correlation between the log $IC_{50}$ in WT and KO cells for each compound. Drugs with a higher level of toxicity for WT (blue) or KO (red) cells are highlighted. **B)** Western blot showing expression of Akt and housekeeping controls (actin, Lamin B1 and H3) in WT and lamin KO A/C cells in different cellular fractions (cytosol, soluble nuclear, and chromatin-bound).
